# Early Tetrapodomorph Biogeography: Controlling for Fossil Record Bias in Macroevolutionary Analyses

**DOI:** 10.1101/726786

**Authors:** Jacob D. Gardner, Kevin Surya, Chris L. Organ

## Abstract

The fossil record provides direct empirical data for understanding macroevolutionary patterns and processes. Inherent biases in the fossil record are well known to confound analyses of this data. Sampling bias proxies have been used as covariates in regression models to test for such biases. Proxies, such as formation count, are associated with paleobiodiversity, but are insufficient for explaining species dispersal owing to a lack of geographic context. Here, we develop a sampling bias proxy that incorporates geographic information and test it with a case study on early tetrapodomorph biogeography. We use recently-developed Bayesian phylogeographic models and a new supertree of early tetrapodomorphs to estimate dispersal rates and ancestral habitat locations. We find strong evidence that geographic sampling bias explains supposed radiations in dispersal rate (potential adaptive radiations). Our study highlights the necessity of accounting for geographic sampling bias in macroevolutionary and phylogenetic analyses and provides an approach to test for its effect.

## 1. Introduction

Our understanding of macroevolutionary patterns and processes are fundamentally based on fossils. The most direct evidence for taxonomic origination and extinction rates come from the rock record, as do evidence for novelty and climate change unseen in data sets gleaned from extant sources. There are no perfect data sets in science; there are inherent limitations and biases in the rock record that must be addressed when we form and test paleobiological hypotheses. For instance, observed stratigraphic ranges of fossils can mislead inferences about diversification and extinction rates (Raup and Boyajian, 1988; Signor and Lipps, 1982). Observed species diversity is also known to increase with time due to the preferential preservation and recovery of fossils in younger geological strata—referred to as “the Pull of the Recent” (Jablonski et al., 2003). Large and long-surviving clades with high rates of early diversification tend to result in an illusionary rate slow-down as diversification rates revert back to a mean value—referred to as “the Push of the Past” (Budd and Mann, 2018). Paleobiologists test and account for these biases when analyzing diversification and extinction at local and global scales (Alroy et al., 2001; Benson et al., 2010; Benson and Butler, 2011; Benson and Upchurch, 2013; Benton et al., 2013; Foote, 2003; Jablonski et al., 2003; Koch, 1978; Lloyd, 2012; Sakamoto et al., 2016a, 2016b). These bias-detection and correction techniques include fossil occurrence subsampling (Alroy et al., 2001; Jablonski et al., 2003; Lloyd, 2012); correcting origination, extinction, and sampling rates using evolutionary predictive models (Foote, 2003); the use of residuals from diversity-sampling models (Benson et al., 2010; Benson and Upchurch, 2013; Sakamoto et al., 2016b); and the incorporation of sampling bias proxies as covariates in regression models (Benson et al., 2010; Benson and Butler, 2011; Benton et al., 2013; Sakamoto et al., 2016a). Benton et al. (2013), studying sampling bias proxies, demonstrated that diversity through time closely tracks formation count (Benton et al., 2013). However, case studies in England and Wales suggest that proxies for terrestrial sedimentary rock volume (such as formation count) do not accurately explain paleobiodiversity, particularly if the fossil record is patchy (Dunhill et al., 2014a, 2014b, 2013). Marine outcrop area and paleoecological-associated facies changes are, however, associated with shifts in paleobiodiversity (Dunhill et al., 2014b, 2013). Moreover, Benton et al. (2013) argue that the direction of causality between paleobiodiversity and formation count is unclear; there may be a common cause to explain their covariation, such as sea level (Benton et al., 2013). Nonetheless, formation count is a widely-used sampling bias proxy in phylogenetic analyses of macroevolution (O’Donovan et al., 2018; Sakamoto et al., 2016a, 2016b; Tennant et al., 2016a, 2016b). The advent of computational modeling approaches, particularly phylogenetic comparative methods, has made it easier to include proxies, like formation count, into models. Additional sampling bias proxies used in these studies include occurrence count, valid taxon count, and specimen completeness and preservation scores. Absent from these proxies is geographic context, which could confound many types of macroevolutionary analyses.

Despite advancements made in understanding the origin and evolution of early tetrapodomorphs, biogeographical studies are hindered by the incompleteness of the early tetrapodomorph fossil record. For example, “Romer’s Gap” represents a lack of tetrapodomorph fossils from the end-Devonian to mid-Mississippian, a period crucial for understanding early tetrapodomorph diversification. Recent collection efforts recovered tetrapodomorph specimens from “Romer’s Gap”, suggesting that a collection and preservation bias explains this gap (Clack et al., 2017; Marshall et al., 2019). In addition, a trackway site in Poland demonstrates the existence of digit-bearing tetrapodomorphs 10 million years before the earliest elpistostegalian body fossil, showcasing the limitation of body fossils to reveal evolutionary history (Niedźwiedzki et al., 2010). A recent study by Long et al. (2018) leveraged phylogenetic reconstruction of early tetrapodomorphs to frame hypotheses about the origin of major clades, as well as their dispersal patterns, including the hypothesis that stem-tetrapodomorphs dispersed from Eastern Gondwana to Euramerica. However, this study did not use phylogenetic comparative methods to estimate ancestral geographic locations or to model dispersal patterns.

Here, we present a phylogeographic analysis of early tetrapodomorphs. Our goals are: 1) to construct a phylogenetic supertree of early tetrapodomorphs that synthesizes previous phylogenetic reconstructions; 2) to estimate the paleogeographic locations of major early tetrapodomorph clades using recently-developed phylogeographic models that account for the curvature of the Earth; and 3) to test for the influence of geographic sampling bias on dispersal rates. Our results indicate that geographic sampling bias substantially confounds analyses of dispersal and paleogeography. We conclude with a discussion about the necessity of controlling for fossil record biases in macroevolutionary analyses.

## 2. Materials and Methods

### 2.1. Nomenclature

Tetrapoda has been informally defined historically to include all terrestrial vertebrates with limbs and digits (Laurin, 1998). Gauthier et al. (1989) first articulated a phylogenetic definition of Tetrapoda as the clade including the last common ancestor of amniotes and lissamphibians. This definition excludes stem-tetrapodomorphs, like *Acanthostega* and *Ichthyostega*. Stegocephalia was coined by E.D. Cope in 1868 (Cope, 1868), but was more recently used to describe fossil taxa more closely related to tetrapods than other sarcopterygians. A recent cladistic redefinition of Stegocephalia includes all vertebrates more closely related to temnospondyls than *Panderichthys* (Laurin, 1998). Here, we use the definitions of Laurin (1998) for a monophyletic Stegocephalia and of Gauthier et al. (1989) for Tetrapoda, which refers specifically to the crown group. We use Tetrapodomorpha to refer to all taxa closer to the tetrapod crown-group than the lungfish crown-group (Ahlberg, 1998). We additionally use Elpistostegalia (= Panderichthyida) to refer to the common ancestor of all stegocephalians and *Panderichthys* as well as Eotetrapodiformes to refer to the common ancestor of all tristichopterids, elpistostegalians, and tetrapods (Coates and Friedman, 2010).

### 2.2. Supertree

We inferred a supertree of 69 early tetrapodomorph taxa from five edited, published morphological data matrices, focusing on tetrapodomorphs whose previously inferred phylogenetic position bracket the water-land transition (Clack et al., 2017; Friedman et al., 2007; Pardo et al., 2017; Swartz, 2012; Zhu et al., 2017). Since downstream analyses might be sensitive to unequal sample sizes between taxa pre- and post-water-land transition, we did not include several crownward stem-tetrapodomorphs from the original matrices (see Supplementary Material). For each matrix, we generated a posterior distribution of phylogenetic trees using MrBayes 3.2.6 (>Ronquist et al., 2012b). In each case, we ran two Markov chain Monte Carlo (MCMC) replicates for 20,000,000 generations with 25% burn-in, each with four chains and a sampling frequency of 1,000. We used one partition, except for Clack et al.’s (2017) matrix, which was explicitly divided into cranial and postcranial characters. To time-calibrate the trees, we constrained the root ages and employed a tip-dating approach (Ronquist et al., 2012a). Tip dates (last occurrence) were acquired from the Paleobiology Database (PBDB; https://paleobiodb.org/) and the literature (see Supplementary Table 2). Root calibrations (minimum and soft maximum age estimates) were collected from the PBDB and Benton et al. (2015). We also used the fossilized birth-death model as the branch length prior (Didier et al., 2017, 2012; Didier and Laurin, 2018; Gavryushkina et al., 2014; Heath et al., 2014; Stadler, 2010; Zhang et al., 2016). All pairs of MCMC replicates converged as demonstrated by low average standard deviation of split frequencies (<0.005; Lakner et al., 2008; see Supplementary Table 3).

Next, we used the five maximum clade credibility trees (source trees; Supplementary Fig. 1-10) to compute a distance supermatrix using SDM 2.1 (Criscuolo et al., 2006). We then inferred an unweighted neighbor-joining tree (UNJ by Gascuel, 1997) from the distance supermatrix using PhyD* 1.1 (Criscuolo and Gascuel, 2008). The UNJ* algorithm is preferable for matrices based on morphological characters. Unlike most supertree methods, the SDM-PhyD* combination produces a supertree with branch lengths. We rooted the supertree using phytools 0.6.60 (Revell, 2012) by adding an arbitrary branch length of 0.00001 to break the trichotomy at the basal-most node in R 3.5.2 (R Core Team, 2018), designating the dipnomorph *Glyptolepis* as the outgroup.

We qualitatively compared the supertree topology with the published source trees and Marjanović and Laurin’s (2019) Paleozoic limbed vertebrate topologies. We also calculated normalized Robinson-Foulds (nRF) distances (Robinson and Foulds, 1981) using phangorn 2.4.0 (Schliep, 2011) in R to assess the congruency of topologies. In each comparison, polytomies in the supertree or the source tree were resolved in all possible ways using phytools. We then calculated all nRF distances and took an average (see Supplementary Table 4). The supplementary materials include a more detailed description of this approach.

### 2.3. Phylogeography

We obtained paleocoordinate data (paleolatitude and paleolongitude) for 63 early tetrapodomorphs from the PBDB using the GPlates software setting (https://gws.gplates.org/). By default, GPlates estimates paleocoordinates from the midpoint of each taxon’s age range. For 16 taxa that did not have direct paleocoordinate data in the PBDB, we searched for the geological formations and geographic regions within the time range from which they are known and averaged the paleolocations across each valid taxonomic occurrence in the PBDB. If the paleolocation of the formation was not listed in the PBDB, we used published geographic locations of the formations. This level of precision is adequate for world-wide phylogeographic analyses, such as conducted here. Present-day coordinates for these geographic locations were obtained from Google Earth and matched with PBDB entries that date within each taxon’s age range (see Supplementary Table 5). Four additional taxa, *Kenichthys, Koilops, Ossirarus*, and *Tungsenia*, had occurrences in the PBDB but the GPlates software could not estimate their paleocoordinates. For *Koilops* and *Ossirarus*, we used all tetrapodomorph occurrences from the Ballagan Formation of Scotland, UK—a formation in which these two taxa are found (Clack et al., 2017). For *Kenichthys* and *Tungsenia*, we calculated paleocoordinate data from the GPlates website directly using the present-day coordinates from the PBDB (https://gws.gplates.org/#recon-p). This approach did not work for the 16 previously mentioned taxa (see Supplementary Table 5). We therefore obtained paleocoordinate data from nearby entries in the PBDB. We excluded the following taxa from our analyses due to the lack of data and comparable entries in the PBDB: *Jarvikina, Koharalepis, Spodichthys*, and *Tinirau.* We excluded the outgroup taxon, *Glyptolepis*, in our analysis to focus on the dispersal trends within early Tetrapodomorpha. We also excluded *Eusthenodon* and *Strepsodus* because their high estimated dispersal rates—being reported from multiple continents—masked other rate variation throughout the phylogeny and inhibited our downstream analyses from converging on a stable likelihood. We do, however, discuss their geographic implications in Section 4.

A model that incorporates phylogeny is crucial for paleobiogeographic reconstruction because it accounts for both species relationships and the amount of evolutionary divergence (branch lengths). Using continuous paleocoordinate data, rather than discretely-coded regions, allows dispersal trends to be estimated at finer resolutions. Discretely-coded geographic regions also limit ancestral states to the same regions inhabited by descendant species. However, standard phylogenetic comparative methods for continuous data assume a flat Earth because they do not account for spherically structured coordinates (i.e., the proximity of −179° and 179° longitudes). Recently-developed phylogenetic comparative methods for modeling continuous paleocoordinate data, implemented as the ‘geo’ model in the program BayesTraits V3, overcome this hurdle by “evolving” continuous coordinate data on the surface of a globe (O’Donovan et al., 2018). The model is implemented with a Bayesian reversible jump MCMC algorithm to estimate rates of geographic dispersal and ancestral paleolocations simultaneously. To account for the spheroid shape of the globe, the ‘geo’ model converts latitude and longitude data into three-dimensional coordinates while prohibiting moves that penetrate the inside of the globe. Ancestral states, which are converted back to standard latitude and longitude, are estimated for each node of the phylogeny. The method includes a variable rates model to estimate variation in dispersal rate (Venditti et al., 2011). The ‘geo’ model makes no assumptions about the location of geographic barriers or coastlines, but a study on dinosaur biogeography found 99.2% of mean ancestral state reconstructions to be located within the bounds of landmasses specific to the time at which they occurred (O’Donovan et al., 2018). We ran three replicate independent analyses using the Bayesian phylogenetic ‘geo’ model for 100 million iterations each with a 25% burn-in and sampling every 1,000 iterations. We estimated log marginal likelihoods using the Stepping Stone algorithm with 250 stones sampling every 1,000 iterations (Xie et al., 2011). We used Bayes factors (BF) to test whether a variable rates model explained the data better than a uniform rates model. Bayes factors greater than two are considered good evidence in support of the model with the greater log marginal likelihood. We compared estimated rate scalars and ancestral states among the three independent variable rates analyses to check for consistency in our results. Rates of dispersal were estimated for each branch by dividing the average rate scalars by the original branch lengths (scaled by time). We assessed the MCMC convergence of all analyses using Tracer 1.7 (Rambaut et al., 2018).

To test for the effect of sampling bias on dispersal rates, we developed a sampling bias proxy that incorporates geographic context: regional-level formation count. Formation counts are meant to capture multiple biases: uneven global rock exposure, uneven fossil collection and database efforts, and global variation in sediment deposition in environments conducive to preservation. Stage-level (stage-specific) formation count represents the mean number of formations, or distinct rock units, globally known to produce relevant fossils along each terminal branch of a phylogeny. Following the protocol of Sakamoto et al. (2016) and O’Donovan et al. (2018), stage-level formation counts are calculated by taking the average number of formations known from each geological age across the globe that encompass the time period between the taxon’s tip date and its preceding node. These average stage-level formation counts are weighted by the proportion that each terminal branch length covers each geological age. For example, if a terminal branch covers two geological ages (e.g., Frasnian and Famennian) at 30% and 70%, respectively, then the stage-level formation counts from each geological age are weighted by those proportions and then divided by the number of geological ages covered:

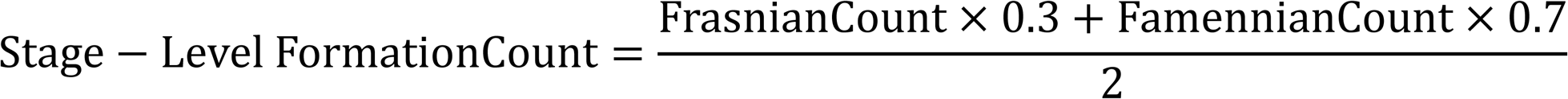

Stage-level formation count is not informed by geography; it is a global metric. It is therefore an inadequate proxy if bias has a strong geographic component (e.g., if the majority of formations recorded are from a specific region or if few formations are exposed within a region). The number of fossil-bearing geological formations, accounting for geographic distribution, is expected to be an important confounding bias in the fossil record. We developed a proxy that includes geographic sampling bias. Our approach breaks down stage-level formation count by geographic region. To account for the arrangement of the continents during the Devonian, Carboniferous, and Permian, we recognized five major regions: Northern Euramerica (including Northeastern Eurasia and Central Asia), Southern Euramerica (North America, Greenland, and Western Europe), Western Gondwana (South America and Africa), Eastern Gondwana (Antarctica, Australia, and Southern Asia), and East Asia (e.g., China). For each branch in the phylogeny, we used the average ancestral state and taxon paleolocation estimates to determine if the branch crossed multiple geographic regions. The number of formations within this time window are totaled for every region covered by the branch and then divided by the number of regions covered. For example, if ancestral state estimates at node 1 and 2 are located in Eastern Gondwana and Southern Euramerica, respectively, then the number of formations recorded in Eastern Gondwana, Southern Euramerica, and the regions in between (i.e., Western Gondwana or Northern Euramerica + East Asia) are counted for that geological age; this total is then divided by the number of geographic regions covered by the entire branch (three for the Western Gondwana route and four for the Northern Euramerica + East Asia route). If the dispersal path between two consecutive ancestral states does not cross any of the five regions, then the number of formations in the inhabited region is counted alone. Figure 1 illustrates an example of how this proxy is measured. This results in the average number of formations present along the dispersal path (at geographic region scale) for each branch in the phylogeny. As with stage-level formation counts, the regional-level formation counts are weighted by the proportion that the branch length covers each geological age. We hypothesize that dispersal rate will inversely correlate with regional-level formation count because we expect that the lack of formations in intermediate regions will lead to inflated dispersal rates. The ‘geo’ model will increase the dispersal rate along a branch to account for the geographic variation observed when there is a lack of intermediate geographic fossil occurrences. This hypothesis can be falsified if high dispersal rates are associated with larger average numbers of formations along dispersal paths. Benton et al. (2013) provide a global sample of tetrapod-bearing rock formations known for each geological age from the Middle Devonian through the Triassic. We supplemented these lists with stratigraphic units known to produce sarcopterygian fossils entered in the PBDB (collected on December 10^th^, 2018).

**1.**
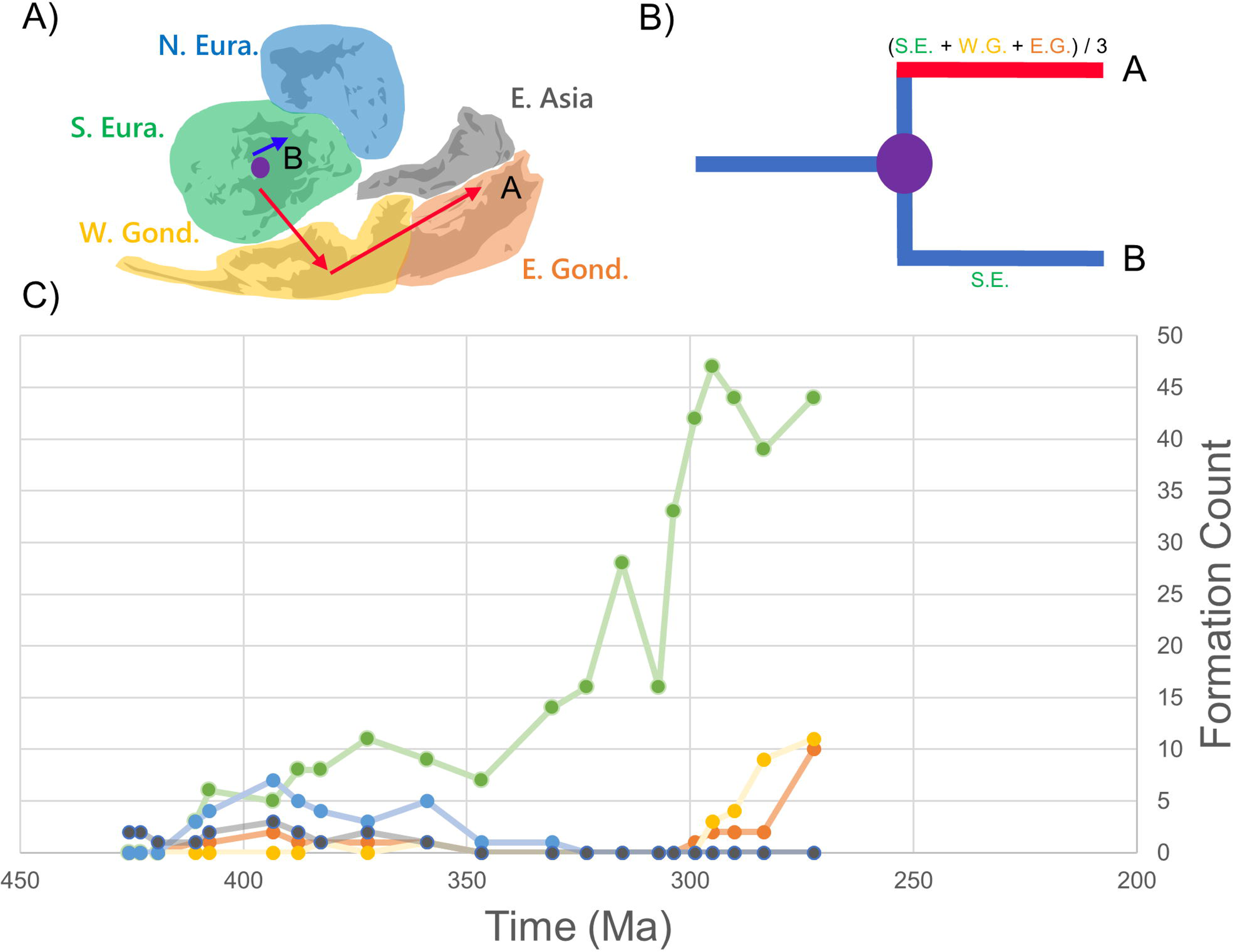
Example of how the regional-level formation count proxy is calculated. A) Five major geographic regions are highlighted by color in the Devonian map. Red arrows represent a branch-specific dispersal path to species A, beginning in Southern Euramerica and ending in Eastern Gondwana. The blue arrow represents the dispersal path to species B. B) The phylogeny of species A and B scaled by time, with equal branch lengths to both species, and colored to represent the rate of dispersal (red is fast, blue is slow). For every branch of the tree, the number of formations is counted for every region and for each geological age covered by the dispersal pathway. It is then weighted by the number of geological ages and geographic regions covered. Under the Western Gondwana route scenario, the branch to species A covers three geographic regions, while the branch to species B only covers one. Assuming both branches cover only one geological age, the high dispersal rate for species A can be explained by the lack of recorded geological formations in Western Gondwana. C) A line plot of the formation counts through time, colored by geographic region according to the Devonian map above, shows temporal and geographic variability.

To test for the effect of regional-level formation count bias on dispersal rate, we conducted a non-parametric two-sample, upper-tailed Mann-Whitney *U*-test using the base package ‘stats’ in R (R Core Team, 2018). This approach ranks all branches of the phylogeny by their regional-level formation count and tests if the branches with lower dispersal rates rank higher on average than branches with higher rates. We define “high” vs “low” dispersal rates based on whether or not they are two standard deviations greater than the average rate across the tree. Due to the vast difference in sample size between the two groups (“high rates”: n = 9, “low rates”: n = 111), we bootstrapped the regional-level formation counts from each group with 100,000 replicates. From this bootstrap analysis, we obtained a 95% confidence interval for the summed ranks of the branches with low dispersal rates (n = 100,000 *U*-statistic values). The expected *U*-statistic is 499.5 given the null hypothesis that only 50% of the regional-level formation counts along branches with low rates rank higher than the formation counts with high rates 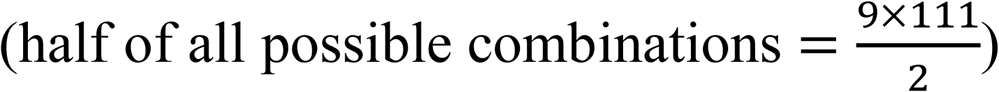. A 95% confidence interval of bootstrapped *U*-statistics that does not include the null expected *U*-statistic is considered good evidence for higher mean dispersal rates along branches with lower regional-level formation counts. The full dataset and code for the phylogeographic analyses can be requested by email to the corresponding author.

Estimated ancestral states do not identify specific dispersal routes, so we conducted sensitivity analyses to test if the dispersal route chosen for counting formations influenced our results. We conceived of three scenarios for dispersal routes between Eastern Gondwana and Southern Euramerica or vice versa: 1) a dispersal route through Western Gondwana; 2) a route through Northern Euramerica and East Asia; and 3) a direct route between Eastern Gondwana and Southern Euramerica. For the first scenario, we averaged the number of formations found in Eastern and Western Gondwana and Southern Euramerica for a given time period. The second scenario is similar to the first but included formation counts from Northern Euramerica and East Asia in place of Western Gondwana. The third scenario only averaged formation counts from Eastern Gondwana and Southern Euramerica.

## 3. Results

### 3.1. Supertree

Topological differences resulted among our supertree, the published source trees, and Marjanović and Laurin’s (2019) tree (Figure 2). In our tree, a polyphyletic “Megalichthyiformes” is the basal-most tetrapodomorph group instead of Rhizodontida (Swartz, 2012; Zhu et al., 2017). Canowindrids and rhizodontids formed an unexpected sister clade to Eotetrapodiformes. Clack et al.’s (2017) five Tournaisian tetrapod taxa cluster together. Colosteidae is rootward of *Crassigyrinus. Caerorhachis* is next to Baphetidae. Baphetidae moved crownward compared to previous topologies (likely because of a small character sample size [Marjanović and Laurin, 2019]). Two crownward nodes are unresolved (polytomous). We retained *Tungsenia* and *Kenichthys* as the oldest and second oldest tetrapodomorphs. Tristichopteridae, Elpistostegalia, Stegocephalia, Aïstopoda, Whatcheeriidae, Colosteidae, Anthracosauria, Dendrerpetidae, and Baphetidae remain monophyletic. Aïstopoda (*Lethiscus* and *Coloraderpeton*) fell rootward to Tetrapoda as reported in Pardo et al. (2017; 2018). The average nRF distances quantify differences in topology (see Supplementary Table 4). On average, there are 39.7% different or missing bipartitions in the source trees compared to the supertree.

**2.**
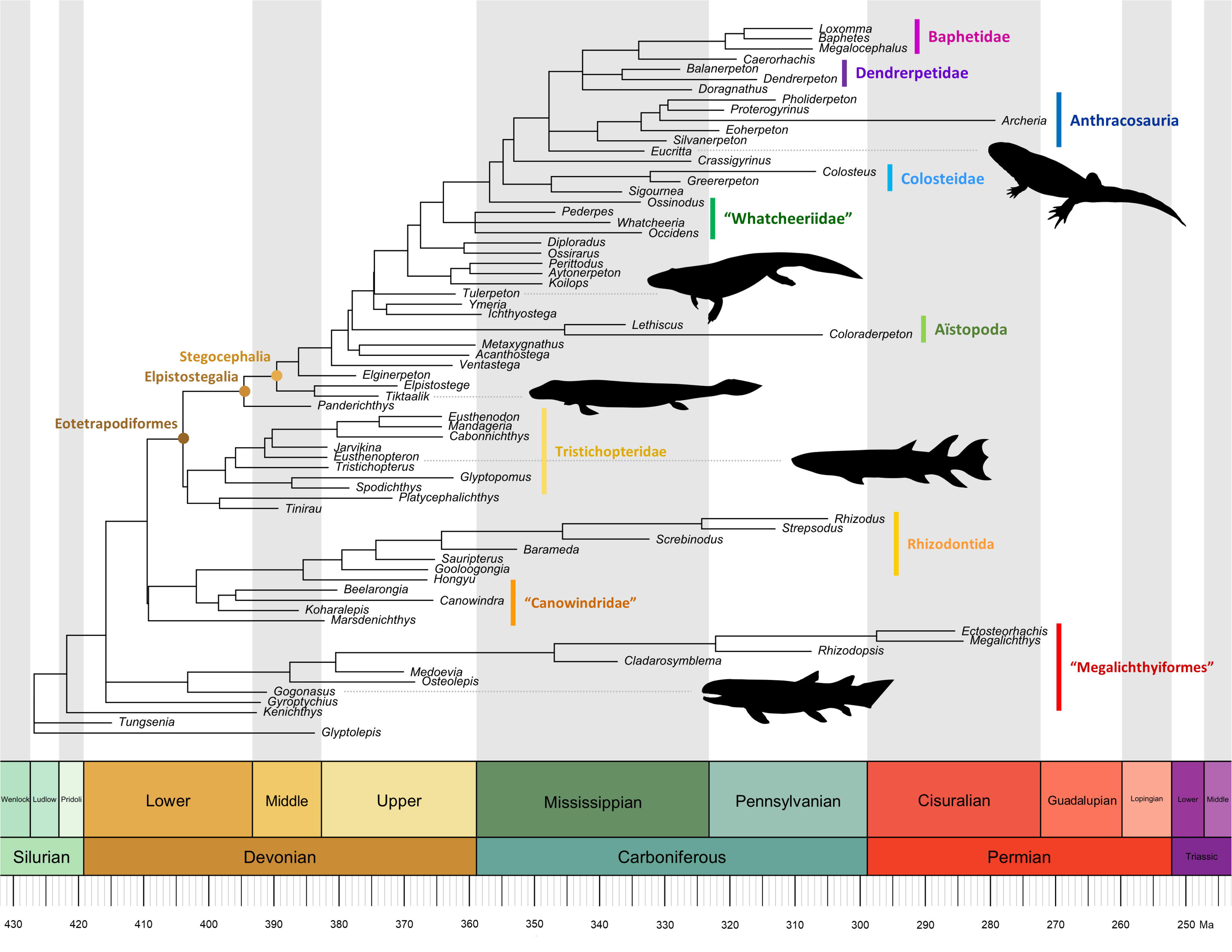
The time-scaled tetrapodomorph supertree. Taxonomic groups in quotes are not monophyletic. Here, *Glyptolepis*, a dipnomorph, is the outgroup. We downloaded the silhouettes from phylopic.org: *Eucritta* and *Greererpeton* by Dmitry Bogdanov (vectorized by Michael Keesey), *Eusthenopteron* by Steve Coombs (vectorized by Michael Keesey), and *Gogonasus* and *Tiktaalik* by Nobu Tamura (CC BY-SA 3.0).

### 3.2. Phylogeography

We found overwhelming support for a variable rates model of geographic dispersal in early tetrapodomorphs (BF = 632.3; Figure 3). The estimated rates across the three replicate runs are consistent (out of 122 branches, only three had a median rate scalar with an absolute value difference among the three runs greater than 3). All rate shifts that were two standard deviations greater than the average dispersal rate were reconstructed dispersal events moving from East Asia to Southern Euramerica, from Eastern Gondwana to Southern Euramerica, or Southern Euramerica to Eastern Gondwana. The fastest estimated dispersal rate occurs along the branch leading to Eotetrapodiformes, moving from Eastern Gondwana to Southern Euramerica (14.34x the average rate). As Long et al. (2018) suggest, we find evidence for an East Asian origin for Tetrapodomorpha but with moderate uncertainty (average estimate ± standard deviation of posterior distribution; longitude_avg_ = 81.5° ± 10.1°, latitude_avg_ = −6.4° ± 8.5°). We also reconstruct an origin for “Megalichthyiformes” that borderlines East Asia and Eastern Gondwana (longitude_avg_ = 107.2° ± 14.1°, latitude_avg_ = −22.6° ± 8.7°), along with an Eastern Gondwana origin for the clade uniting “Canowindridae” and Rhizodontida (longitude_avg_ = 137.1° ± 8.2°, latitude_avg_ = −32.0° ± 4.7°). We recover a Southern Euramerican origin for Eotetrapodiformes, consistent with previous studies (longitude_avg_ = −12.5° ± 7.0°, latitude_avg_ = −19.4° ± 6.4°). A Southern Euramerican origin was also found for Tristichopteridae (longitude_avg_ = −12.7° ± 6.9°, latitude_avg_ = −19.7° ± 6.3°) and Elpistostegalia (longitude_avg_ = −12.3° ± 5.5°, latitude_avg_ = −13.5° ± 5.3°). As expected in a phylogenetic comparative analysis, uncertainty in estimated node states increases toward the root. However, despite the level of uncertainty within a single run, only three nodes have mean ancestral state values that are greater than an absolute value of 5° among the replicate three runs.

**3.**
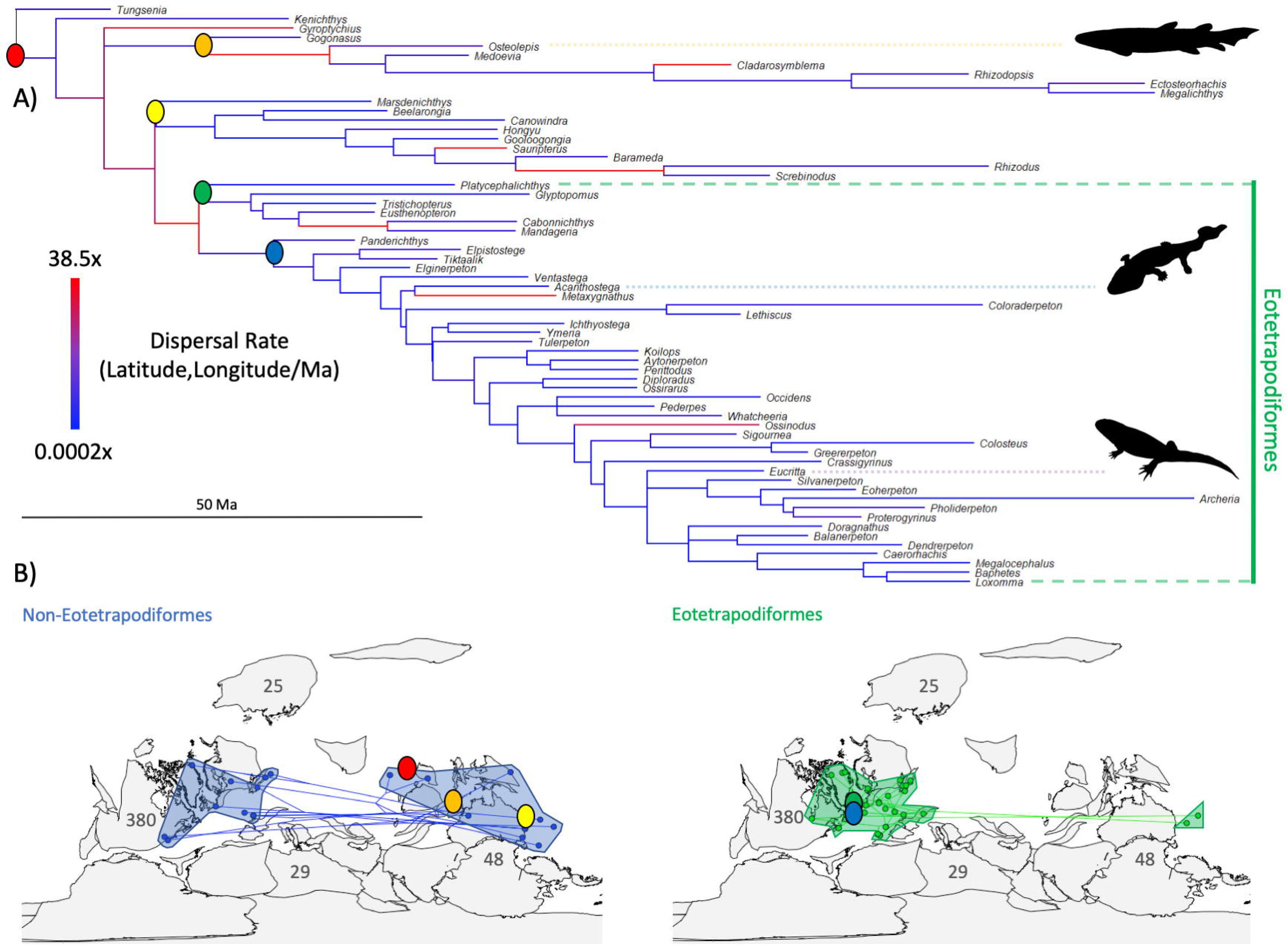
A) Trimmed tetrapodomorph phylogeny with mapped rates of dispersal. Cooler (bluish) colors represent slower rates and warmer (reddish) colors represent faster rates. B) Non-eotetrapodiform (left in blue) and eotetrapodiform (right in green) trees and taxon paleolocations plotted on a map of the Middle Devonian. Transparent polygons illustrate broad geographic regions of sampled taxa in Southern Euramerica, Eastern Gondwana, and East Asia. Numbers show the total number of geological formations recorded from each major geographic region (Eastern Gondwana and East Asia combined). Colored circles show average paleolocations of major clades estimated by the ‘geo’ model and indicated in the tree above. Red circle: Tetrapodomorpha, orange: “Megalichthyiformes”, yellow: “Canowindridae” + Rhizodontidae, green: Tristichopteridae, and blue: Elpistostegalia. Phylogeny with mapped dispersal rates was produced in BayesTrees (http://www.evolution.rdg.ac.uk/BayesTrees.html). Middle Devonian tree and paleolocation plots were made using the ‘phylo-to-map’ function in the R package, phytools (Revell, 2012). Middle Devonian map was sourced from the R package, paleoMap (Rothkugel and Varela, 2015). Tetrapodomorph silhouettes were sourced from phylopic.org: *Eucritta* by Dmitry Bogdanov (vectorized by T. Michael Keesey), *Osteolepis* by Nobu Tamura, and *Acanthostega* by Mateus Zica.

We find good evidence that geographic sampling bias influences dispersal rate estimates, regardless of the route used (95% CI: Western Gondwana route *U* = [800, 928]; Northern Euramerica + East Asia route *U* = [832, 946]; direct route *U* = [729, 889]; no scenario includes the null *U* = 499.5; Figure 4 and Supplementary Figures 12-13). A *U*-statistic considerably higher than 499.5 suggests that branches with high dispersal rates have lower regional-level formation counts, on average, than branches with low rates. One can also interpret the null *U*-statistic of 499.5 as a 50% probability that a random branch with a low dispersal rate will rank higher in its regional-level formation count than a random branch with a high dispersal rate. With bootstrapping, we are 95% confident that the probability of a random branch with a low dispersal rate having a higher regional-level formation count than a random branch with a high rate is 72.97–88.99% for the more conservative ‘direct route’ scenario. Under the more liberal ‘Northern Euramerica + East Asia route’ scenario, the probabilities are 83.28–94.69%. In sum, branches with high dispersal rates (two standard deviations greater than average) have a smaller number of recorded formations, on average, along their reconstructed dispersal path.

**4.**
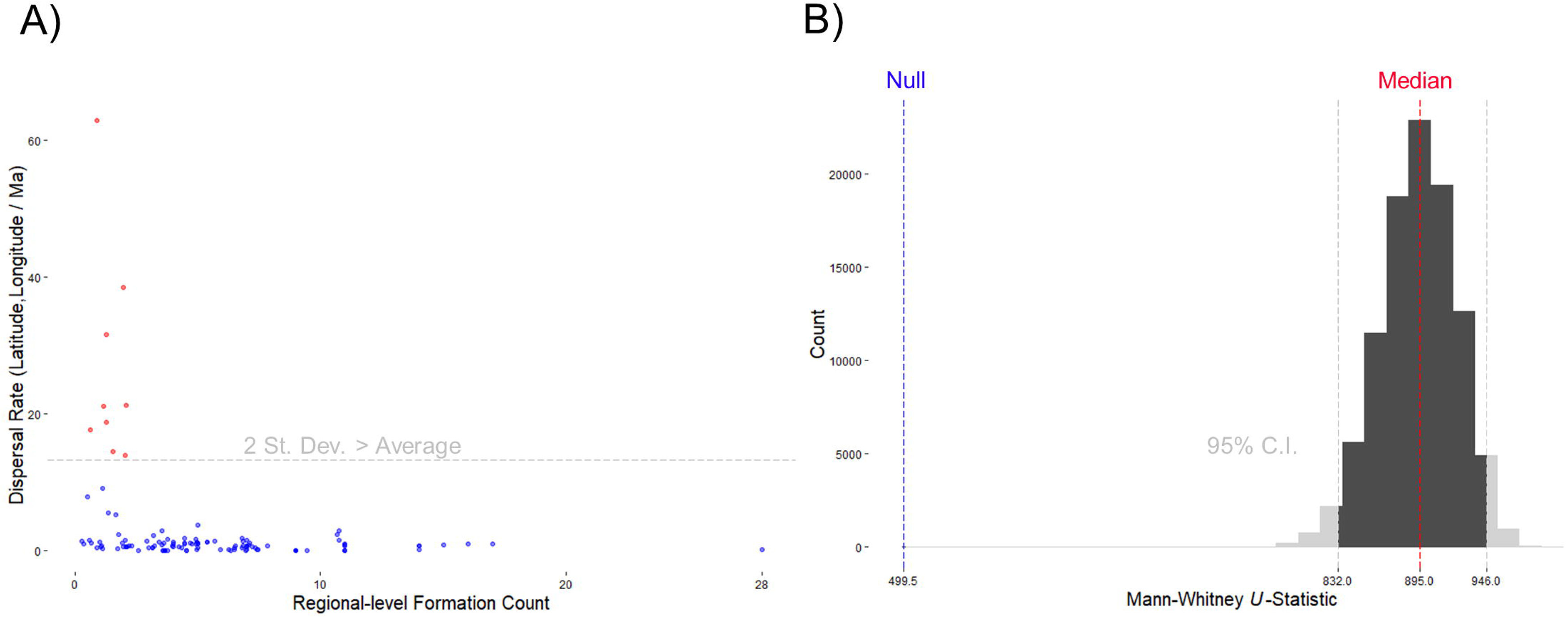
A) Scatter-plot of the average dispersal rates over the regional-level formation counts for each branch of the phylogeny, using the Northern Euramerica + East Asia route scenario. Points colored by the dispersal rate being above (red) or below (blue) two standard deviations greater than the average rate across the tree. B) Histogram of the bootstrapped *U*-statistics. Values outside of the 95% confidence interval are grayed out. The median and null expected *U*-statistics are indicated by the red and blue dotted lines, respectively. The null expected *U*-statistic is based on the null hypothesis that 50% of the branches with low dispersal rates will have a greater regional-level formation count than branches with higher rates. Rejecting the null hypothesis suggests that estimated dispersal rates are biased and correlate with regional-level formation count.

Our results cannot be explained by a fossil record that is more complete through time (Pull of the Recent). A regression model relating regional-level formation count to the minimum age of each branch shows only a weak relationship (slope = -0.044, r^2^ = 0.1, *P* < 0.001). However, total global (stage-level) formation count (which does not account for geographic variation) does show potential bias from Pull of the Recent (slope = −0.3, r^2^ = 0.71, *P* < 0.0001). If dispersal rates are biased by the increase in number of formations globally, we would also expect to see elevated dispersal rates decrease toward the tips, but a regression model relating stretched branch lengths with time is not supported (slope = −0.025, r^2^ = 0.006, *P* = 0.41).

## 4. Discussion

We expected to infer high dispersal rates for closely related taxa that are distributed across the globe. Our results, unadjusted for geographic bias in the fossil record, confirm this notion. However, we also find a compelling statistical association between high dispersal rates and a low number of formations along dispersal paths—a patchy fossil record is driving inferences of high dispersal rates. Although we did not test for a correlation between dispersal rate and previously used proxies, such as valid taxon count and stage-level formation count, these proxies do not offer clear predictions for explaining dispersal rate variation. High dispersal rate variation is inferred when closely related taxa are geographically separate. For example, valid taxon count cannot explain geographic rate variation because spatial information is lacking in this bias proxy and because sister taxa are likely to have similar counts (these data are phylogenetically structured). Stage-level formation counts will also not explain dispersal rate variation, particularly if high rate variation exists within the same geological age. Assuming geological formations are evenly exposed and sampled worldwide, low stage-level formation counts should yield geographically variable fossil species and, therefore, drive high dispersal rate variation. However, formations are not evenly exposed or recorded in geological/paleontological databases, including the PBDB. Our formation count table demonstrates this bias (Table 1). Without geographic context, stage-level formation count cannot distinguish between global and local regions. For example, the geological ages that have the highest recorded number of formations are restricted to Southern Euramerica where the majority of eotetrapodiform taxa have been discovered. The association between high formation counts in specific regions and high paleobiodiversity in those regions is likely not a coincidence and has a clear impact on how we interpret dispersal history. The earliest tetrapodomorphs are known from China and Australia at geological ages where relatively few formations are recorded outside of East Asia and Eastern Gondwana. The basal-most ancestral state estimates reconstruct paleolocations in East Asia (not surprisingly). This inference (hypothesis) is predicated on the lack of geological formations recorded outside of East Asia during this time period. In addition, the majority of more crownward taxa and their reconstructed ancestral states are located in North America and Europe at geological ages in which relatively fewer formations are known elsewhere. This bias may heavily influence any conclusions made on the location and habitat of the tetrapod water-land transition. Recently discovered taxa could help mitigate this problem by increasing the power of taxon sampling (Heath et al., 2014), such as *Tutusius* and *Umzantsia* from South Africa (Gess and Ahlberg, 2018). However, the current lack of cladistic coding for these taxa excludes them from phylogeny-based analyses. The taxonomic resolution of globally-occurring species, like *Eusthenodon* and *Spodichthys*, also impacts current models of species dispersal history because of their relatively uniform distribution (Long et al., 2018). *Eusthenodon* and *Spodichthys* represent possible cases where taxonomic resolution is too coarse for phylogeographic analyses. Including these species inhibited our MCMC algorithms from reaching convergence. Widely distributed cosmopolitan species that lack intermediate geographic occurrences increase the uncertainty of parameter estimates within phylogeographic models, as is the case here for these two species.

**Table 1:** Regional- and stage-level (total) formation counts through time.

| Period | Epoch | Age | End Time (Ma) | Northern Euramerica | Southern Euramerica | Western Gondwana | Eastern Gondwana | East Asia | Total |
| --- | --- | --- | --- | --- | --- | --- | --- | --- | --- |
| Permian | Cisuralian | Kungurian | 272.95 | 10 | 44 | 11 | 0 | 0 | 65 |
| Permian | Cisuralian | Artinskian | 283.5 | 2 | 39 | 9 | 0 | 0 | 50 |
| Permian | Cisuralian | Sakmarian | 290.1 | 2 | 44 | 4 | 0 | 0 | 50 |
| Permian | Cisuralian | Asselian | 295 | 2 | 47 | 3 | 0 | 0 | 52 |
| Pennsylvanian | Late | Gzhelian | 298.9 | 1 | 42 | 0 | 0 | 0 | 43 |
| Pennsylvanian | Late | Kasimovian | 303.7 | 0 | 33 | 0 | 0 | 0 | 33 |
| Pennsylvanian | Middle | Moscovian | 307 | 0 | 16 | 0 | 0 | 0 | 16 |
| Pennsylvanian | Early | Bashkirian | 315.2 | 0 | 28 | 0 | 0 | 0 | 28 |
| Mississippian | Late | Serpukhovian | 323.2 | 0 | 16 | 0 | 0 | 0 | 16 |
| Mississippian | Middle | Viséan | 330.9 | 0 | 14 | 0 | 1 | 0 | 15 |
| Mississippian | Early | Tournaisian | 346.7 | 0 | 7 | 0 | 1 | 0 | 8 |
| Devonian | Late | Famennian | 358.9 | 1 | 9 | 1 | 5 | 1 | 17 |
| Devonian | Late | Frasnian | 372.2 | 1 | 11 | 0 | 3 | 2 | 17 |
| Devonian | Middle | Givetian | 382.7 | 1 | 8 | 1 | 4 | 1 | 15 |
| Devonian | Middle | Eifelian | 387.7 | 1 | 8 | 0 | 5 | 2 | 16 |
| Devonian | Early | Emsian | 393.3 | 2 | 5 | 0 | 7 | 3 | 17 |
| Devonian | Early | Pragian | 407.6 | 1 | 6 | 0 | 4 | 2 | 13 |
| Devonian | Early | Lochkovian | 410.8 | 1 | 3 | 0 | 3 | 1 | 8 |
| Silurian | Prídolí | Prídolí | 419.2 | 0 | 0 | 0 | 0 | 1 | 1 |
| Silurian | Ludlow | Ludfordian | 423 | 0 | 0 | 0 | 0 | 2 | 2 |
| Silurian | Ludlow | Gorstian | 425.6 | 0 | 0 | 0 | 0 | 2 | 2 |

Phylogenetic studies on macroevolution also often fail to incorporate data from the fossil record itself, such as trace fossil occurrences. Non-anatomical data often contribute to our understanding of taxonomic originations, including chiridian (or digit-possessing) tetrapodomorphs for which trace fossil evidence exists about 10 million years before the first elpistostegalian body fossils (Niedźwiedzki et al., 2010). The inclusion of additional data from trace fossils could radically alter our current models of species dispersal history. Finally, it is important to note that the sampling bias proxies are also constrained by database curation biases. Phylogenetic studies on macroevolutionary trends now regularly leverage public databases, such as the PBDB, which allows larger and broader studies. It is unclear how patchy entries, on taxonomic occurrences and geological formations, for example, interact with other biases inherent in the fossil record. Caution is therefore warranted when these databases are mined, as is the case here.

## 5. Conclusions

Phylogenetic studies on macroevolution have not previously incorporated geographic context, which could influence a wide variety of analyses. We demonstrate here that phylogeographic methods are influenced by geographic sampling variability. We develop a simple sampling bias proxy that incorporates geographic information and show that it explains variation in estimated dispersal rates. The majority of elevated dispersal rates are associated with large-scale movements between major landmasses that have very few, if any, relevant geological formations in between. Our analysis is also unlikely to be influenced by “Pull of the Recent”-like effects. Although not the first supertree for early tetrapodomorphs (Ruta et al., 2003), this study presents the first (to our knowledge) with branch lengths, making it useable for phylogenetic comparative analyses. The new supertree comprises many of the major clades previously inferred, but also recovers new ones that will be subject to scrutiny in future studies (discussed further in the Supplementary Material). This supertree should be useful to researchers who aim to use phylogenetic comparative methods to test hypotheses on the evolution of early tetrapodomorphs. In sum, our study estimates ancestral geographical reconstructions consistent with previously hypothesized dispersal patterns in early tetrapodomorphs. We also find that rates of dispersal are strongly influenced by geographic sampling bias. We suggest that researchers incorporate this proxy in phylogeny-based macroevolutionary studies that could be influenced by spatial distribution of the fossil record.

## Acknowledgements

We thank the MSU Macroevolution Lab, Jack Wilson, and Matt Lavin for helpful discussions, as well as David Marjanović and John Long for their helpful reviews. We also thank Nathalie Bardet, Eric Buffetaut, Annelise Folie, Emmanuel Gheerbrant, Alexandra Houssaye, and Michel Laurin for the invitation to contribute to this special issue celebrating the life and accomplishments of Jean-Claude Rage.

