## Supplementary Material for "Early Tetrapodomorph Biogeography: Controlling for Fossil Record Bias in Macroevolutionary Analyses"

### Supplementary Materials

### Table of Contents

1. Supertree
  1.1. Data
    1.1.1. Morphological data matrices 1
    1.1.2. Tip dates 1
    1.1.3. Root calibrations 3
  1.2. Analyses
    1.2.1. Bayesian phylogenetic inference 3
    1.2.2. Distance supermatrix 4
    1.2.3. Unweighted neighbor-joining 4
    1.2.4. Rooting and plotting 5
    1.2.5. Tree comparisons 5
2. Phylogeography
  2.1. Data
    2.1.1. Paleocoordinate locations 6

Supplementary tables
  Table 1: Data matrix summary 1
  Table 2: Tip dates 2
  Table 3: Average standard deviation of split frequencies values 4
  Table 4: Average normalized Robinson-Foulds (nRF) distances 5
  Table 5: Taxon paleolocation sources 6

Supplementary figures
  Figure 1: The phylogenetic tree inferred using Friedman et al.'s (2007) matrix 9
  Figure 2: The phylogenetic tree inferred using Swartz's (2012) matrix 10
  Figure 3: The phylogenetic tree inferred using Clack et al.'s (2017) matrix 11
  Figure 4: The phylogenetic tree inferred using Pardo et al.'s (2017) matrix 12
  Figure 5: The phylogenetic tree inferred using Zhu et al.'s (2017) matrix 13
  Figure 6: The time-scaled phylogenetic tree inferred using Friedman et al.'s (2007) matrix 14
  Figure 7: The time-scaled phylogenetic tree inferred using Swartz's (2012) matrix 15
  Figure 8: The time-scaled phylogenetic tree inferred using Clack et al.'s (2017) matrix 16
  Figure 9: The time-scaled phylogenetic tree inferred using Pardo et al.'s (2017) matrix 17
  Figure 10: The time-scaled phylogenetic tree inferred using Zhu et al.'s (2017) matrix 18
  Figure 11: Supertree's branch confidence values 19
  Figure 12: Formation counts with low dispersal rates tend to rank higher than those with higher rates (Western Gondwana route scenario) 20
  Figure 13: Formation counts with low dispersal rates tend to rank higher than those with higher rates (direct route scenario) 21

Supplementary files description
  1. Supertree
    1.1. Data 22
    1.2. Analysis 22
    1.3. Publication 22
  2. Phylogeography
    2.1. Data 23
    2.2. Analysis 23

### 1. Supertree

#### *1.1. Data*

##### *1.1.1. Morphological data matrices*

To maximize stem-tetrapodomorph taxon sample size, we collected five published data matrices containing unordered, multistate morphological characters (Friedman et al., 2007; Swartz, 2012; Clack et al., 2017; Pardo et al., 2017; Zhu et al., 2017) (see Table 1). Since downstream analyses might be sensitive to unequal sample sizes between taxa pre- and post-water-land transition, we didn't include several crownward stem-tetrapods from the original matrices. All taxa more crownward than *Baphetes* and *Eucritta* in Pardo et al. (2017), except *Balanerpeton* and *Dendrerpeton*, were disregarded. *Asaphestera*, *Casineria*, *Discosauriscus*, *Edops*, *Eryops*, *Gephyrostegus*, *Hyloplesion*, *Microbrachis*, *Paleothyris*, *Seymouria*, and *Westlothiana* were removed from Clack et al.'s (2017) data matrix. Future studies should include more taxa on either end of the water-land transition. Lastly, we chose the supertree approach because it's infeasible to construct a morphological supermatrix. One needs to recode characters and assess redundancies.

| **Source** | **# of taxa** | **# of characters** | **Outgroup** |
|---|---|---|---|
| Friedman et al. (2007) | 13 | 216 | *Glyptolepis* |
| Swartz et al. (2012) | 41 | 204 | *Glyptolepis* |
| Clack et al. (2017) | 33 | 213 | *Eusthenopteron* |
| Pardo et al. (2017) | 18 | 370 | *Eusthenopteron* |
| Zhu et al. (2017) | 33 | 169 | *Glyptolepis* |

**Table 1.** Data matrix summary. Number of taxa refers to the number of stem-tetrapodomorphs included in our analyses. For three data matrices, we chose *Glyptolepis* as the outgroup in subsequent Bayesian phylogenetic inferences because it seemed to be the skeletally most complete dipnomorph.

We made several corrections to the data matrices. We changed *Ymeria's* state in character 343 of Pardo et al.'s (2017) matrix from 3 to a question mark (?) because that character only has two states. In Clack et al.'s (2017) matrix, we changed the states of *Silvanerpeton* in character 18, *Proterogyrinus* in character 31, and *Pholiderpeton* in character 66 from 2, 3, and 2, respectively, to question marks (?). These character states exceed the maximum number of states. Additionally, we substituted parentheses in Swartz's (2012) and Pardo et al.'s (2017) matrices with curly brackets for consistency.

Here, we must explicitly acknowledge that we didn't guarantee that every specimen used to score character states is skeletally mature. We didn't order characters (see Rineau et al., 2015, 2018). Further, we didn't check correlations between characters (see Guillerme and Brazeau [2018] for discussions regarding this issue). These caveats could have biased phylogenetic inference.

##### *1.1.2. Tip dates*

We collected tip dates (genus-level minimum ages) from the Paleobiology Database (PBDB; https://paleobiodb.org/). If minimum age data were unavailable from the PBDB, we checked the localities where researchers found the genus, chose the youngest one and collected the upper boundary of the locality's age based on http://fossilworks.org (see Table 2).

| Taxon | Tip date (Ma) | Geochronologic unit | Reference |
|---|---|---|---|
| *Acanthostega* | 358.90 | Late Famennian | PBDB |
| *Archeria* | 279.30 | Middle Kungurian | PBDB |
| *Aytonerpeton* | 346.70 | Late Tournaisian | PBDB |
| *Balanerpeton* | 326.40 | Middle Serpukhovian | PBDB |
| *Baphetes* | 307.00 | Late Moscovian | PBDB |
| *Barameda* | 346.70 | Late Tournaisian | PBDB |
| *Beelarongia* | 376.10 | Frasnian | Long, (1987) |
| *Cabonnichthys* | 360.70 | Famennian | Ahlberg and Johanson, (1997) |
| *Caerorhachis* | 318.10 | Middle Bashkirian | PBDB |
| *Canowindra* | 360.70 | Late Devonian | Thomson, (1973) |
| *Cladarosymblema* | 326.40 | Viséan | Fox et al., (1995) |
| *Coloraderpeton* | 307.00 | Late Moscovian | PBDB |
| *Colosteus* | 306.95 | Early Kasimovian | PBDB |
| *Crassigyrinus* | 323.20 | Late Serpukhovian | PBDB |
| *Dendrerpeton* | 314.60 | Early Moscovian | PBDB |
| *Diploradus* | 346.70 | Late Tournaisian | PBDB |
| *Doragnathus* | 323.20 | Late Serpukhovian | PBDB |
| *Ectosteorhachis* | 272.30 | Early Roadian | PBDB |
| *Elginerpeton* | 376.10 | Middle Frasnian | PBDB |
| *Elpistostege* | 372.20 | Late Frasnian | PBDB |
| *Eoherpeton* | 318.10 | Middle Bashkirian | PBDB |
| *Eucritta* | 330.90 | Late Viséan | PBDB |
| *Eusthenodon* | 360.70 | Late Famennian | Clement, (2002) |
| *Eusthenopteron* | 372.20 | Late Frasnian | PBDB |
| *Glyptolepis* | 382.40 | Early Frasnian | PBDB |
| *Glyptopomus* | 360.70 | Late Famennian | Lebedev and Lukševičs, (2017) |
| *Gogonasus* | 382.40 | Early Frasnian | Long et al., (2006) |
| *Gooloogongia* | 360.70 | Famennian | Johanson and Ahlberg, (1998) |
| *Greererpeton* | 323.20 | Late Serpukhovian | PBDB |
| *Gyroptychius* | 383.70 | Middle Devonian | Newman et al., (2015) |
| *Hongyu* | 360.70 | Famennian | Zhu et al., (2017) |
| *Ichthyostega* | 358.90 | Late Famennian | PBDB |
| *Jarvikina* | 379.50 | Middle Frasnian | Lebedev et al., (2010) |
| *Kenichthys* | 382.70 | Late Givetian | PBDB |
| *Koharalepis* | 382.40 | Early Frasnian | Young et al., (1992) |
| *Koilops* | 346.70 | Late Tournaisian | PBDB |
| *Lethiscus* | 336.00 | Middle Viséan | PBDB |
| *Loxomma* | 306.95 | Early Kasimovian | PBDB |
| *Mandageria* | 360.70 | Famennian | Johanson and Ahlberg (1997) |
| *Marsdenichthys* | 376.10 | Frasnian | Holland et al., (2010) |
| *Medoevia* | 360.70 | Late Devonian | Lebedev, (1995) |
| *Megalichthys* | 272.30 | Early Roadian | PBDB |
| *Megalocephalus* | 306.95 | Early Kasimovian | PBDB |

| | | | |
|---|---|---|---|
| *Metaxygnathus* | 358.90 | Late Famennian | PBDB |
| *Occidens* | 330.90 | Late Viséan | PBDB |
| *Ossinodus* | 330.90 | Late Viséan | PBDB |
| *Ossirarus* | 346.70 | Late Tournaisian | PBDB |
| *Osteolepis* | 358.90 | Late Famennian | PBDB |
| *Panderichthys* | 382.40 | Early Frasnian | PBDB |
| *Pederpes* | 345.30 | Early Viséan | PBDB |
| *Perittodus* | 346.70 | Late Tournaisian | PBDB |
| *Pholiderpeton* | 311.45 | Middle Moscovian | PBDB |
| *Platycephalichthys* | 360.70 | Late Famennian | Lebedev et al., (2010) |
| *Proterogyrinus* | 318.10 | Middle Bashkirian | PBDB |
| *Rhizodopsis* | 298.90 | Late Gzhelian | PBDB |
| *Rhizodus* | 298.90 | Late Gzhelian | PBDB |
| *Sauripterus* | 358.90 | Late Famennian | PBDB |
| *Screbinodus* | 326.40 | Viséan | Andrews, (1985) |
| *Sigournea* | 336.00 | Middle Viséan | PBDB |
| *Silvanerpeton* | 326.40 | Middle Serpukhovian | PBDB |
| *Spodichthys* | 376.10 | Frasnian | Snitting, (2008) |
| *Strepsodus* | 307.00 | Late Moscovian | PBDB |
| *Tiktaalik* | 372.20 | Late Frasnian | PBDB |
| *Tinirau* | 383.70 | Late Givetian | Swartz, (2012) |
| *Tristichopterus* | 379.50 | Early Frasnian | Bishop, (2013) |
| *Tulerpeton* | 360.70 | Late Famennian | PBDB |
| *Tungsenia* | 407.60 | Late Pragian | PBDB |
| *Ventastega* | 358.90 | Late Famennian | PBDB |
| *Whatcheeria* | 336.00 | Middle Viséan | PBDB |
| *Ymeria* | 358.90 | Late Famennian | PBDB |

**Table 2.** Tip dates.

*1.1.3. Root calibrations*

We collected clade minimum and soft maximum ages from the PBDB and Benton et al. (2015) to calibrate tree roots. For Friedman et al. (2007), Swartz (2012), and Zhu et al. (2017), the least inclusive clade with age estimates is Rhipidistia (minimum age = 408.0 Ma; soft maximum age = 427.9 Ma). For Clack et al. (2017) and Pardo et al. (2017), we used the maximum ages of *Eusthenopteron*, *Panderichthys*, and *Spodichthys* (one of the basalmost taxa in Eotetrapodiformes) as the minimum age of the least inclusive clade (383.7 Ma). And the mean age of the clade is represented by the minimum age of Tetrapodomorpha (407.6 Ma).

*1.2. Analyses*

*1.2.1. Bayesian phylogenetic inference*

For each matrix, we generated a posterior distribution of phylogenetic trees using MrBayes 3.2.6 (Ronquist et al., 2012b). We wanted to have trees with branch length information and to standardize the inference process as much as possible. Here, we included details absent from the manuscript. Aside from using outgroups designated in Table 1, we also constrained the ingroup. For

Clack et al.'s (2017) matrix, we allowed the gamma shape parameter, state frequency, and rate to vary across partitions. Next, we conditioned on coding only variable characters (Lewis Mkv model [Lewis, 2001] corrects for ascertainment bias). Therefore, 150, 8, 6, 103, and 6 constant characters in Friedman et al.'s (2007), Swartz's (2012), Clack et al.'s (2017), Pardo et al.'s (2017), and Zhu et al.'s (2017) matrices, respectively, were ignored. Moreover, we used gamma-shaped rate variation across sites (four categories; Yang, 1994). Harrison and Larsson (2015) found that the four rate category discrete approximation is sufficient to approximate a gamma rate distribution. We used an exponentially-distributed prior for the gamma shape parameter. An exponentially-distributed prior for the gamma shape parameter results in higher marginal likelihoods than a uniformly-distributed prior (Harrison and Larsson, 2015). To allow variable evolutionary rates over time, we used the Independent Gamma Rate (IGR) model (Lepage et al., 2007). As a prior for the morphological clock rate, we used a truncated normal distribution. Further, we used offset exponential priors for root and tree ages and fixed tip dates (see Ronquist et al., 2012a). Although we employed the fossilized birth-death model (FBD; Stadler, 2010; Didier et al., 2012, 2015; Heath et al., 2014; Gavryushkina et al., 2014; Zhang et al., 2016; Didier and Laurin 2018) as a branch length prior, we didn't allow for sampled ancestors.

In each inference, we ran two Markov chain Monte Carlo (MCMC) replicates for 20,000,000 generations, each with four chains, a sampling frequency of 1,000, and a diagnostics frequency of 5,000. MrBayes employs Metropolis-coupled version of the MCMC (Metropolis et al., 1953; Hastings, 1970; Geyer, 1991). We discarded the first 25% samples as burn-in. We also used BEAGLE 2.1 (Ayres et al., 2012) to decrease computational time. Unless specified above, we used the default settings. Finally, we chose to output maximum clade credibility trees (Fig. 1-10).

We diagnosed MCMC convergence between runs using the average standard deviation (SD) of split frequencies (Lakner et al., 2008). The values in all five inferences were less than 0.005 (see Table 3). We also assessed convergence using minimum effective sample size (ESS) and potential scale reduction factor (PSRF by Gelman and Rubin, 1992) values. All these metrics showed that within each inference, runs converged.

| **Inference** | **Average SD of split frequencies** |
| --- | --- |
| Friedman et al. (2007) | 0.002578 |
| Swartz et al. (2012) | 0.004706 |
| Clack et al. (2017) | 0.003893 |
| Pardo et al. (2017) | 0.003184 |
| Zhu et al. (2017) | 0.004766 |

**Table 3.** The average standard deviation of split frequencies values between runs in all inferences were less than 0.005.

*1.2.2. Distance supermatrix*

First, we converted the maximum clade credibility trees (source trees) to Newick files using FigTree 1.4.3 (Rambaut, 2017). Then, we combined all the Newick trees into a single PHYLIP file. We inputted this file to SDM 2.1 (Criscuolo et al., 2006) and computed a distance supermatrix. Trees were weighted using their sizes.

*1.2.3. Unweighted neighbor-joining*

We inferred the supertree from the distance supermatrix using a modified unweighted neighbor-joining (Gascuel, 1997) algorithm (UNJ*) implemented in PhyD* 1.1 (Criscuolo and Gascuel, 2008). We allowed polytomies and only positive branch lengths. Furthermore, we chose to output confidence values at branches (Guénoche and Garreta, 2000), which were suited for incomplete distance matrices. Most values are above 50. However, we didn't understand why our supertree contains branches with zero confidence values (Fig. 11), especially when nearby nodes had high posterior probabilities in the source trees. Unless specified above, we used the default settings.

*1.2.4. Rooting and plotting*

We read the supertree into R 3.5.2 (R Core Team, 2018) using APE 5.2 (Paradis and Schliep, 2019), rooted and saved it using phytools 0.6.60 (Revell, 2012), converted it to a Newick file using FigTree, and converted it again to a .trees file using BayesTreesConverter 1.3 (http://www.evolution.rdg.ac.uk/BayesTrees.html). BayesTraits 3.0.1 (Pagel 1999; http://www.evolution.rdg.ac.uk/BayesTraitsV3.0.1/BayesTraitsV3.0.1.html) can't process a tree with a polytomous root node. So, we added an arbitrary branch length of 0.00001 to break the trichotomy.

Lastly, we plotted the complete supertree using strap 1.4 (Bell and Lloyd, 2015), Cairo 1.5.9 (Urbanek and Horner, 2015), and APE in R. Since *Archeria* has the longest path length, we scaled the tree using *Archeria's* tip date (279.3 Ma) despite *Megalichthys* and *Ectosteorhachis'* tip dates (272.3 Ma).

*1.2.5. Tree comparisons*

We compared the supertree with the published source tree and Marjanović and Laurin's (2019) Paleozoic limbed vertebrate tree regarding topology. Due to small stem-tetrapod sample size, we ignored Friedman et al.'s (2007) topology. Additionally, we prioritized published Bayesian over maximum parsimony trees whenever possible.

Lastly, we compared the supertree topology with the published source tree topologies using normalized Robinson-Foulds (nRF) distances (Robinson and Foulds, 1981) implemented in phangorn 2.4.0 (Schliep, 2011) in R. We first wrote Newick files for the source tree topologies. For Clack et al.'s (2017) Bayesian tree, we designated *Eusthenopteron* as the outgroup. Afterward, we pruned the supertree to match the tips in individual source trees using APE in R. To match Swartz's (2012) tree, we collapsed several clades (Rhizodontidae, Megalichthyidae, Whatcheeridae, Colosteidae, Baphetidae, total-group Lissamphibia, and Embolomeri). There appears to be no actual taxon in the clade "other stem-group amniotes" in Swartz's (2012) tree figure. In each comparison, polytomies in the supertree or the source tree were resolved in all possible ways using phytools. Then, we calculated all possible nRF distances and took an average (see Table 4).

| **Inference** | **Average nRF distance (%)** |
|---|---|
| Friedman et al. (2007) | 25.0 |
| Swartz et al. (2012) | 27.1 |
| Clack et al. (2017) | 67.7 |
| Pardo et al. (2017) | 33.3 |
| Zhu et al. (2017) | 45.6 |
| **Average** | 39.7 |

**Table 4.** The average of average normalized Robinson-Foulds (nRF) distances is 39.7%. Thus, there are, on average, 39.7% different or missing bipartitions in the source trees compared to the supertree.

### 2. Phylogeography

#### *2.1. Data*

##### *2.1.1. Paleocoordinate locations*

We obtained paleocoordinate data (paleolatitude and paleolongitude) for 65 early tetrapodomorphs from the PBDB using the GPlates software setting (https://gws.gplates.org/). For 16 taxa that did not have direct paleocoordinate data in the PBDB, we searched for the geologic formations and geographic regions, while encapsulating the time range, from which they were discovered and averaged all valid tetrapodomorph occurrences from those formations and regions. If not the paleolocation of the formation entry given by the PBDB, we used the closest geographic location from where a publication stated the formation is located. Although not the precise location, a more-general geographic location (e.g. township, county, or country) should suffice for the global scale that we're conducting analyses. Below is a table of the locations we used for each of the 16 taxa.

| Taxon | Paleolocation source | Reference | Notes |
|---|---|---|---|
| *Acanthostega* | PBDB | - | |
| *Archeria* | PBDB | - | |
| *Aytonerpeton* | PBDB | - | |
| *Balanerpeton* | PBDB | - | |
| *Baphetes* | PBDB | - | |
| *Barameda* | PBDB | - | |
| *Beelarongia* | PBDB - Avon River Group | - | |
| *Cabonnichthys* | PBDB - New South Wales | Long et al. (2018) | |
| *Caerorhachis* | PBDB | - | |
| *Canowindra* | PBDB - New South Wales | Long et al. (2018) | |
| *Cladarosymblema* | PBDB - Queensland | Long et al. (2018) | |
| *Coloraderpeton* | PBDB | - | |
| *Colosteus* | PBDB | - | |
| *Crassigyrinus* | PBDB | - | |
| *Dendrerpeton* | PBDB | - | |
| *Diploradus* | PBDB | - | |
| *Doragnathus* | PBDB | - | |
| *Ectosteorhachis* | PBDB | - | |
| *Elginerpeton* | PBDB | - | |
| *Elpistostege* | PBDB | - | |
| *Eoherpeton* | PBDB | - | |
| *Eucritta* | PBDB | - | |
| *Eusthenodon* | PBDB - Celsius Bjerg Group, Tula Region, Evieux Formation, New South Wales, Witpoort Formation | Clement et al. (2009), Long et al. (2018) | Not included: outlier rates |
| *Eusthenopteron* | PBDB | - | |

| | | | |
|---|---|---|---|
| *Glyptolepis* | - | - | Not included: outgroup |
| *Glyptopomus* | PBDB - Latvia | Lebedev and Lukševičs (2017) | |
| *Gogonasus* | PBDB | - | |
| *Gooloogongia* | PBDB - New South Wales | Long et al. (2018) | |
| *Greererpeton* | PBDB | - | |
| *Gyroptychius* | PBDB - Estonia, Scotland | Newman et al. (2015) | |
| *Hongyu* | PBDB - Zhongning | - | |
| *Ichthyostega* | PBDB | - | |
| *Jarvikina* | Russia | Young et al. (2013) | Not included: specific region in Russia unknown |
| *Kenichthys* | portal.gplates | - | Used present-day coordinates from PBDB |
| *Koharalepis* | PBDB - Mount Crean, Antarctica | Long et al. (2018) | Not included: no Antarctica entries in PBDB |
| *Koilops* | PBDB - Ballagan Formation | - | |
| *Lethiscus* | PBDB | - | |
| *Loxomma* | PBDB | - | |
| *Mandageria* | PBDB - New South Wales | Long et al. (2018) | |
| *Marsdenichthys* | PBDB - Mount Howitt, Victoria | Long et al. (2018) | |
| *Medoevia* | PBDB - Latvia | Lebedev (1995) | From Belarus, used Latvia occurrences |
| *Megalichthys* | PBDB | - | |
| *Megalocephalus* | PBDB | - | |
| *Metaxygnathus* | PBDB | - | |
| *Occidens* | PBDB | - | |
| *Ossinodus* | PBDB | - | |
| *Ossirarus* | PBDB - Ballagan Formation | - | |
| *Osteolepis* | PBDB | - | |
| *Panderichthys* | PBDB | - | |
| *Pederpes* | PBDB | - | |
| *Perittodus* | PBDB | - | |
| *Pholiderpeton* | PBDB | - | |
| *Platycephalichthys* | PBDB - Latvia | Boisvert et al. (2008) | |
| *Proterogyrinus* | PBDB | - | |
| *Rhizodopsis* | PBDB | - | |
| *Rhizodus* | PBDB | - | |
| *Sauripterus* | PBDB | - | |
| *Screbinodus* | PBDB - Scotland | Andrews (1985) | |
| *Sigournea* | PBDB | - | |
| *Silvanerpeton* | PBDB | - | |
| *Spodichthys* | East Greenland | Snitting (2008) | Not included: PBDB entries for Greenland are outside of age range |
| *Strepsodus* | PBDB (North American *Strepsodus* entries), PBDB - Queensland | Parker et al. (2005) | Not included: outlier rates |

| | | | |
|---|---|---|---|
| *Tiktaalik* | PBDB | - | |
| *Tinirau* | Eureka County, Nevada | Swartz (2012) | Not included: PBDB entries for Eureka Country, NV, are outside of age range |
| *Tristichopterus* | PBDB - Scotland | - | |
| *Tulerpeton* | PBDB | - | |
| *Tungsenia* | portal.gplates | - | Used present-day coordinates from PBDB |
| *Ventastega* | PBDB | - | |
| *Whatcheeria* | PBDB | - | |
| *Ymeria* | PBDB | - | |

**Table 5.** Taxon Paleolocations.

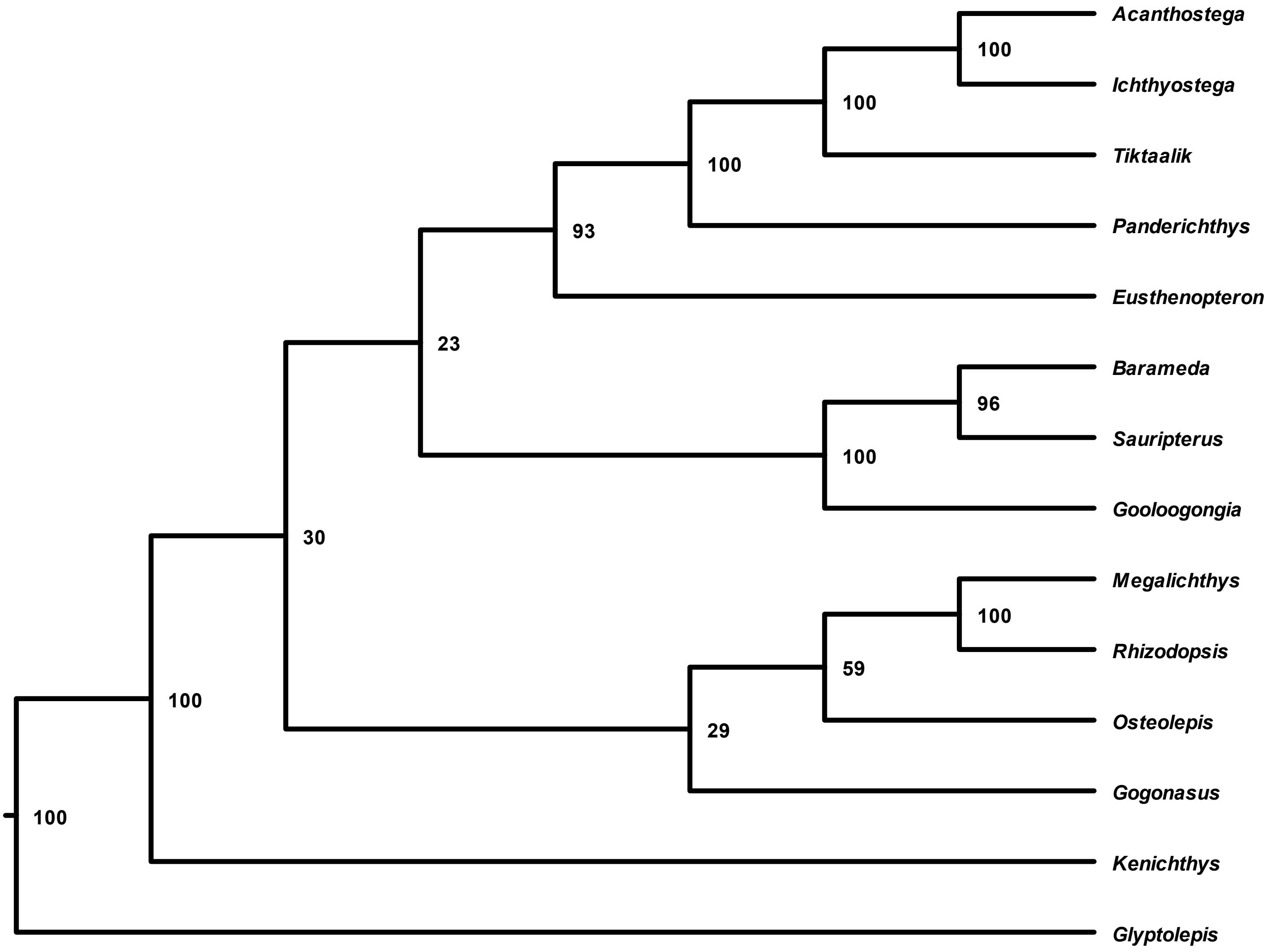


**Fig. 1.** The phylogenetic tree inferred using Friedman et al.'s (2007) matrix. Node values represent percent node posterior probabilities. Note that the hypothesized relationship between Megalichthyiformes, Rhizodontida, and Eotetrapodiformes have low support (30 and 23). We used FigTree to produce this figure.

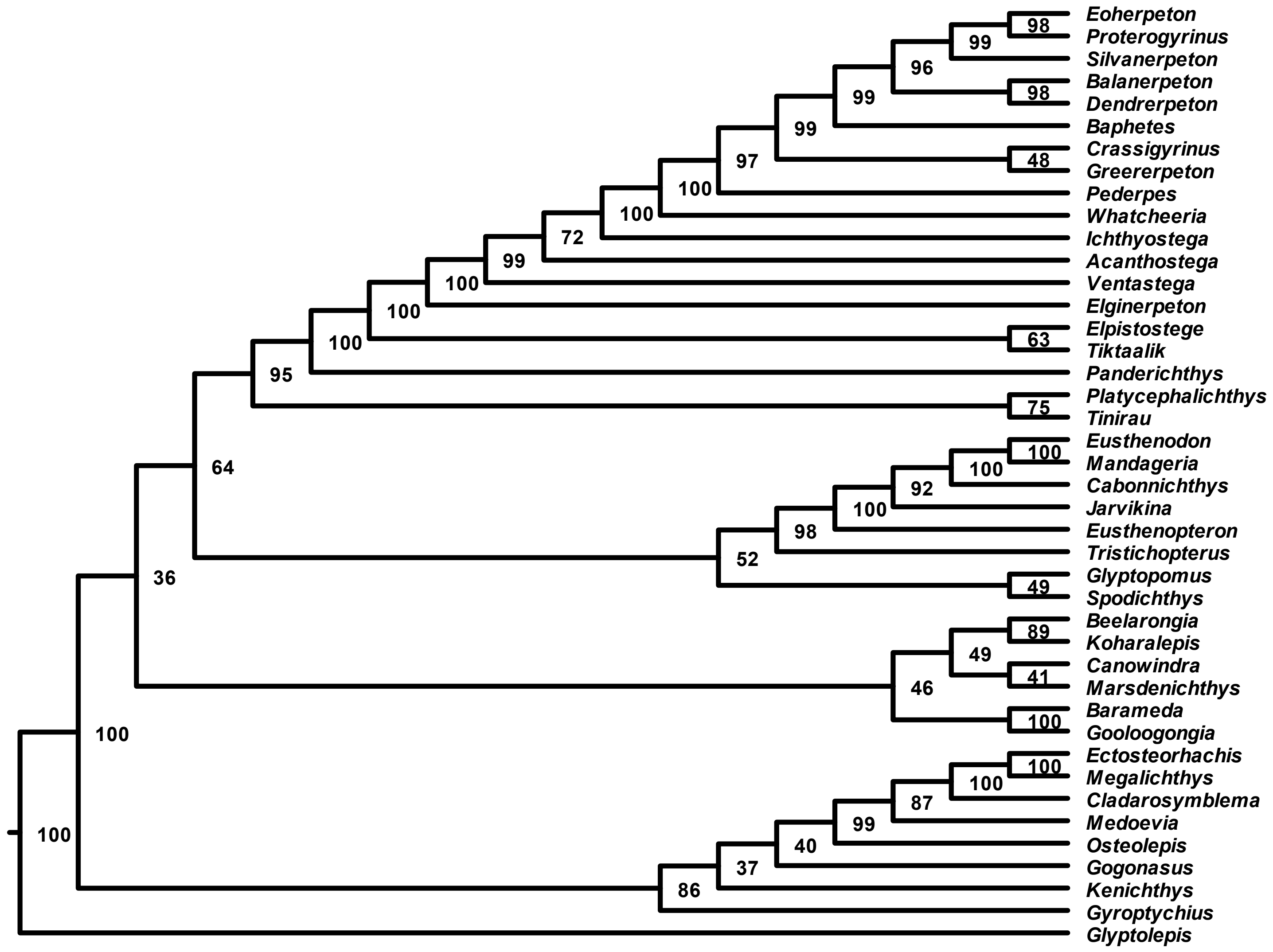


**Fig. 2.** The phylogenetic tree inferred using Swartz's (2012) matrix. Node values represent posterior probabilities (%). The sister taxon relationship between a group of canowindrids and rhizodontids and Eotetrapodiformes has low support (36).

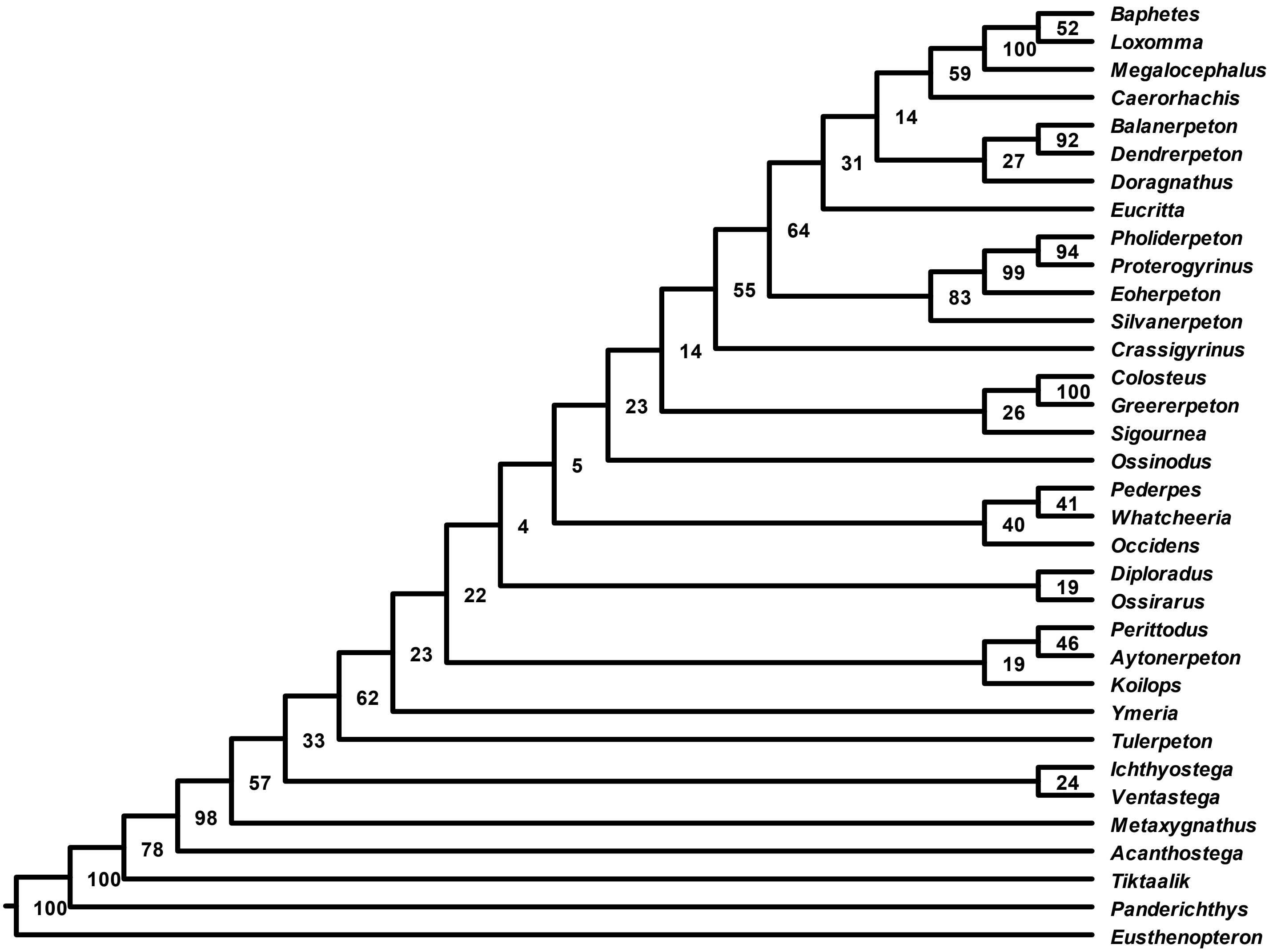


**Fig. 3.** The phylogenetic tree inferred using Clack et al.'s (2017) matrix. Node values represent posterior probabilities (%). Note the low support for multiple backbone nodes.

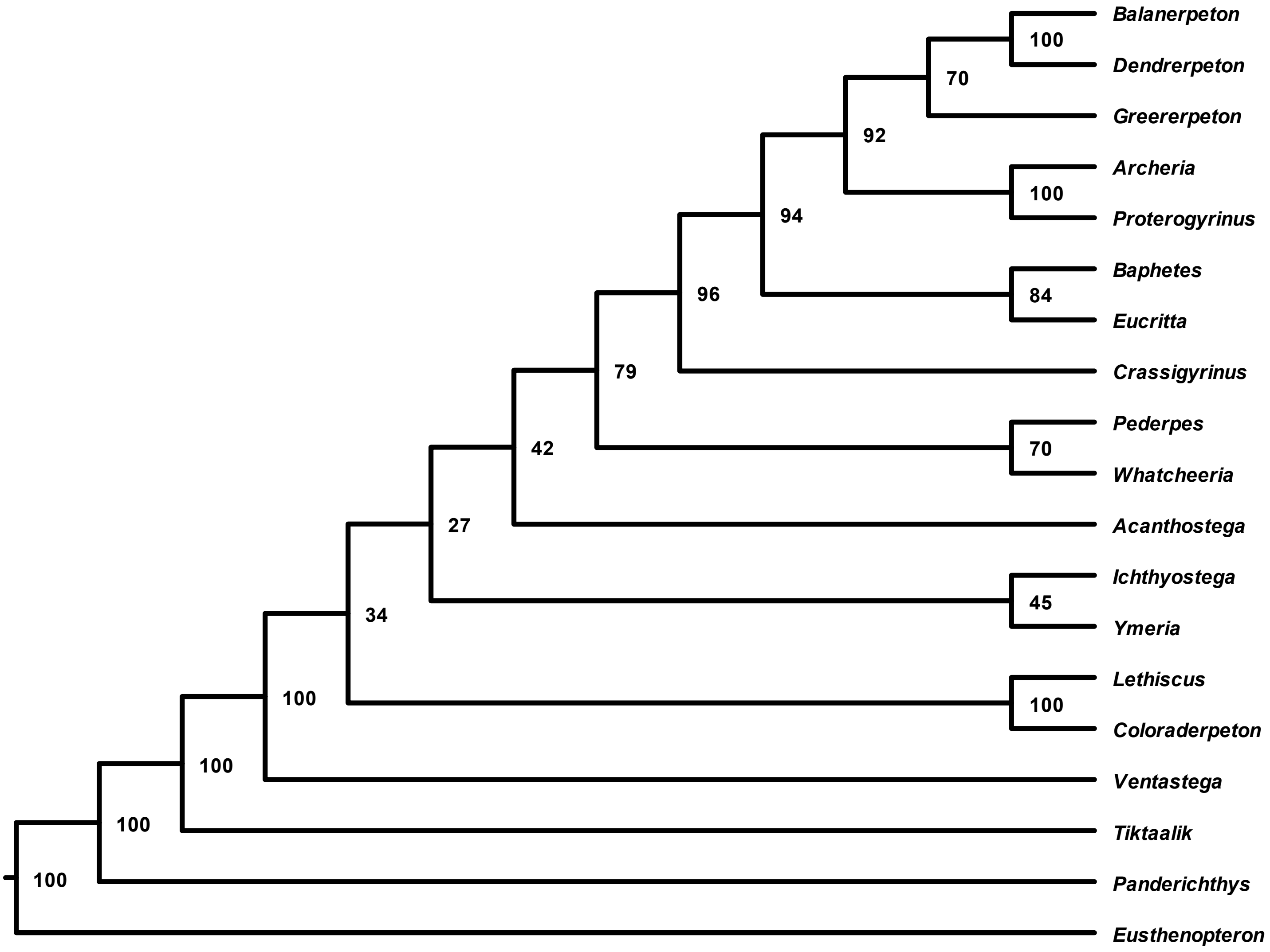


**Fig. 4.** The phylogenetic tree inferred using Pardo et al.'s (2017) matrix. Node values represent posterior probabilities (%). Note the low support for some backbone nodes (34, 27, and 42).

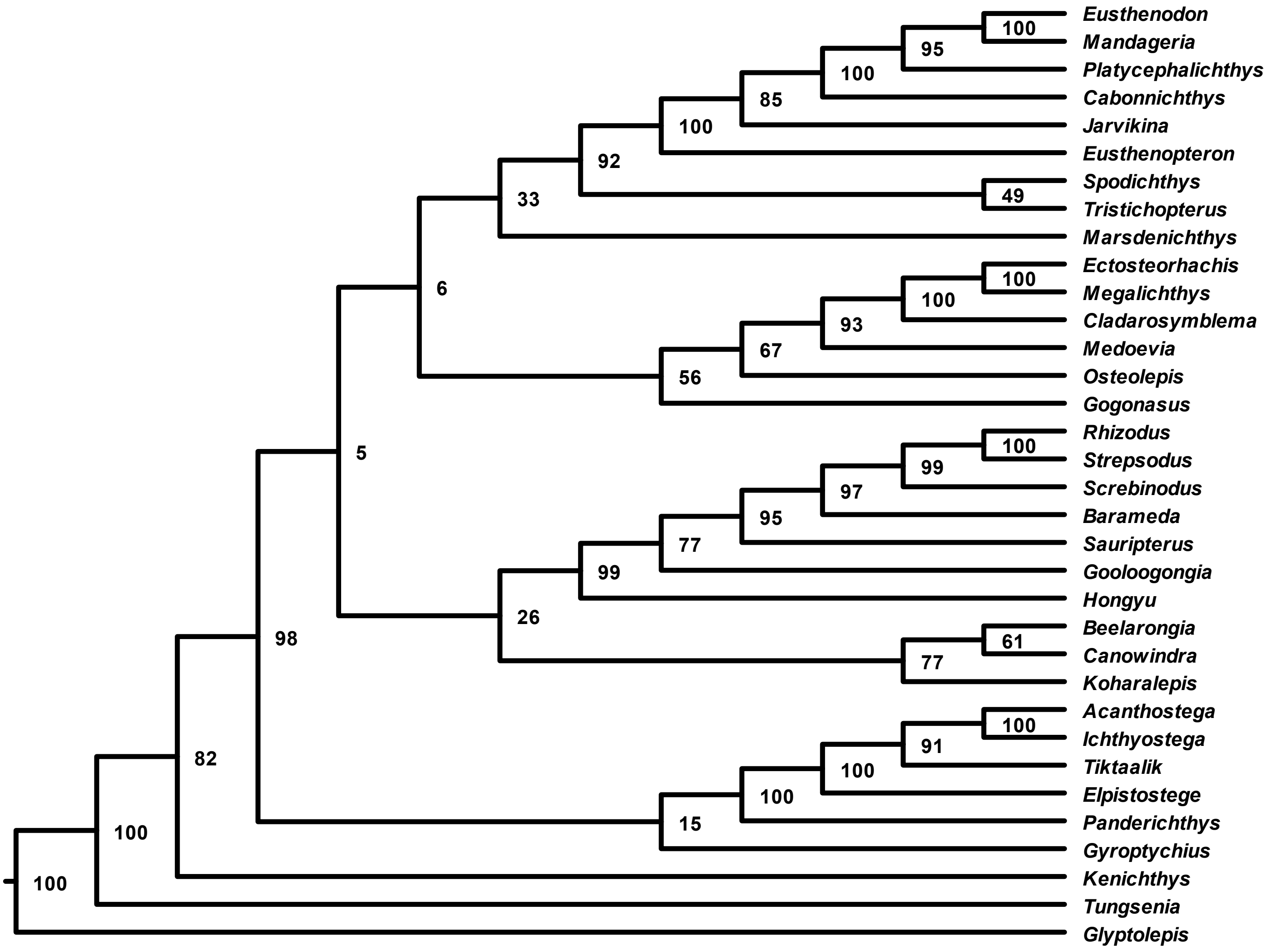


**Fig. 5.** The phylogenetic tree inferred using Zhu et al.'s (2017) matrix. Node values represent posterior probabilities (%). Note that the hypothesized relationship between Canowindridae, Rhizodontida, Megalichthyiformes, and Tristichopteridae have low support (5, 6, and 26). This unconventional topology shows an early divergence of Elpistostegalia from the rest of stem-tetrapods.

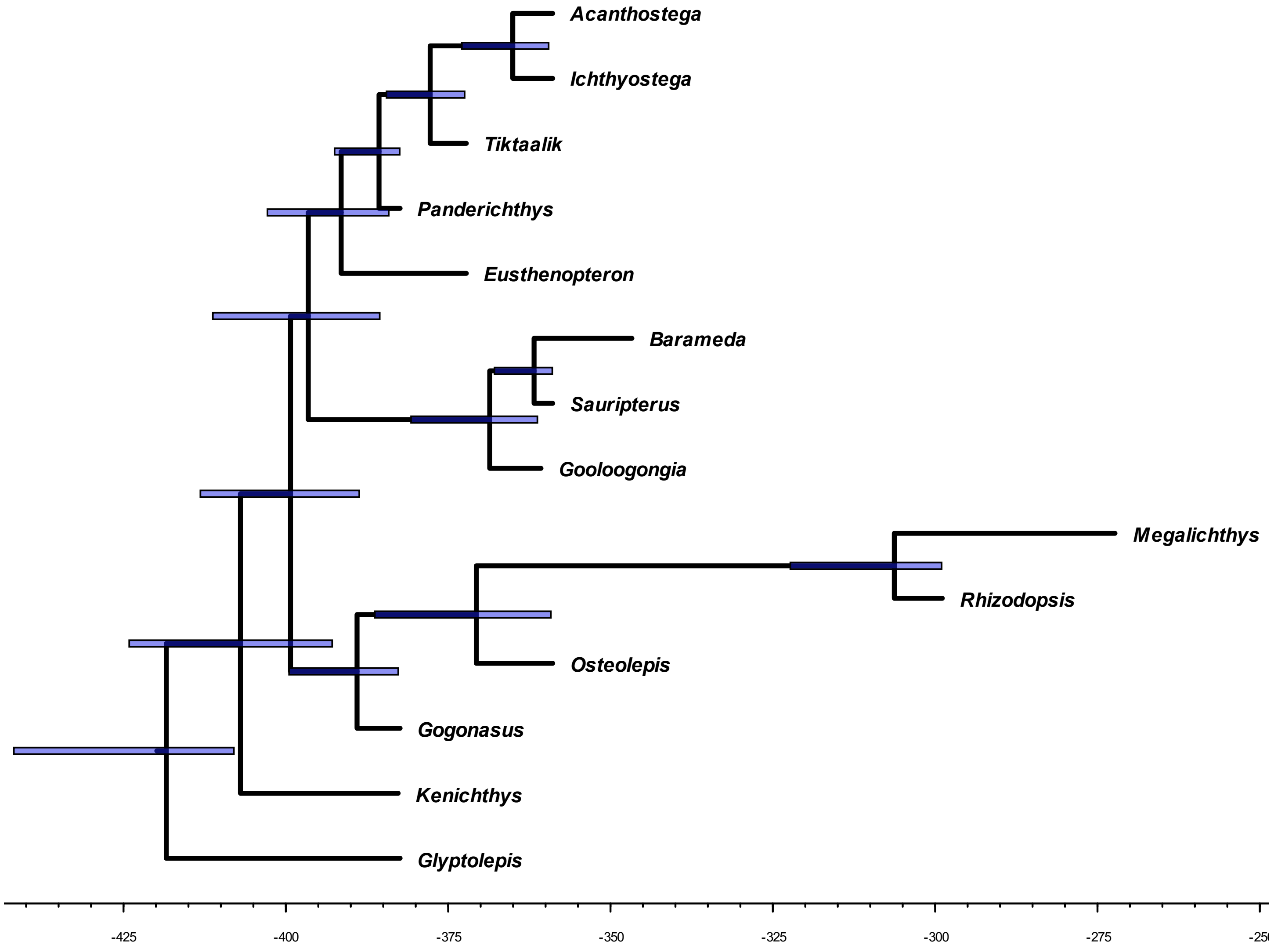


**Fig. 6.** The time-scaled phylogenetic tree inferred using Friedman et al.'s (2007) matrix. Node bars represent 95% highest posterior density (HPD) of node age estimates.

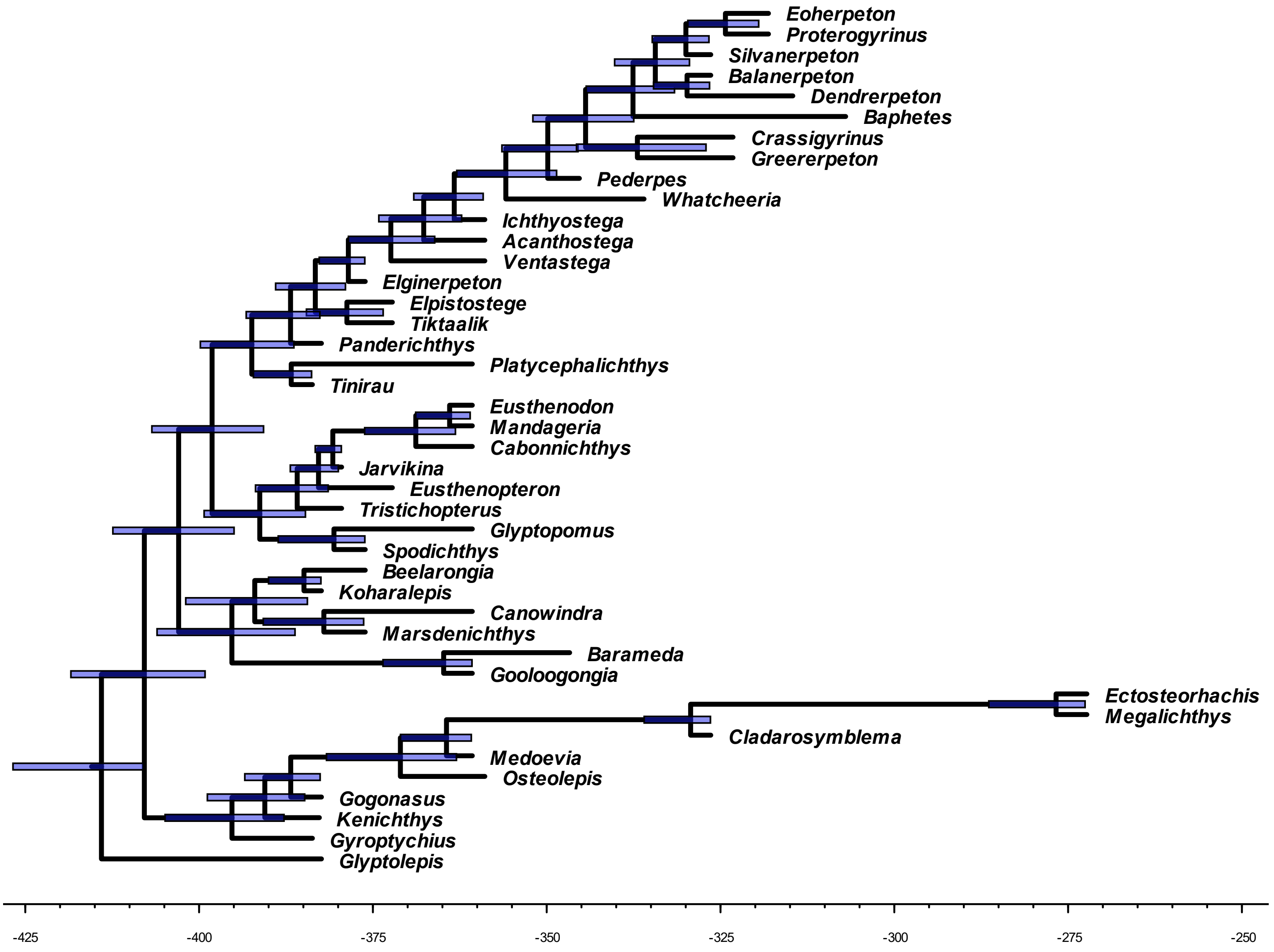


**Fig. 7.** The time-scaled phylogenetic tree inferred using Swartz's (2012) matrix. Node bars represent 95% highest posterior density (HPD) of node age estimates.

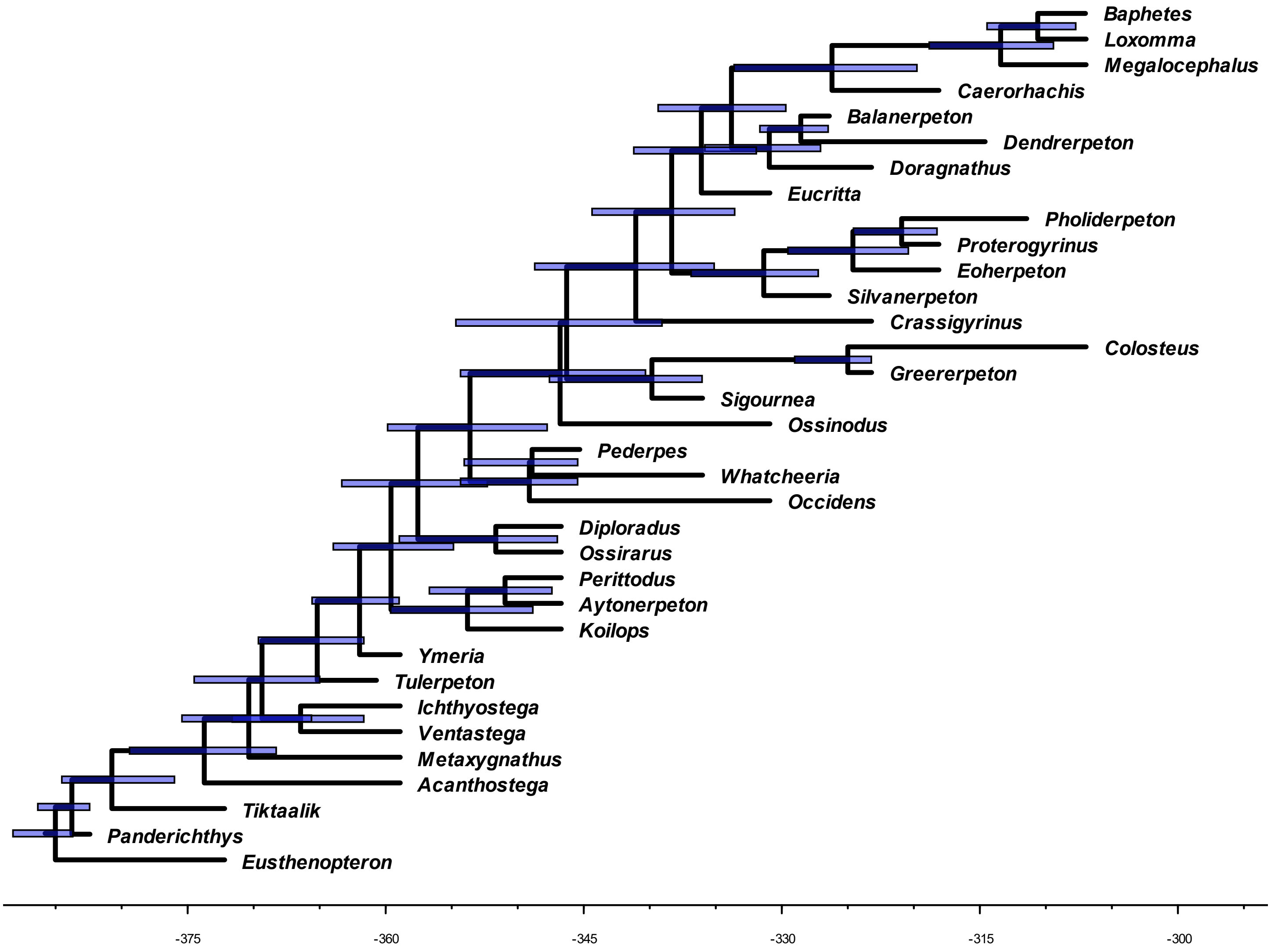


**Fig. 8.** The time-scaled phylogenetic tree inferred using Clack et al.'s (2017) matrix. Node bars represent 95% highest posterior density (HPD) of node age estimates.

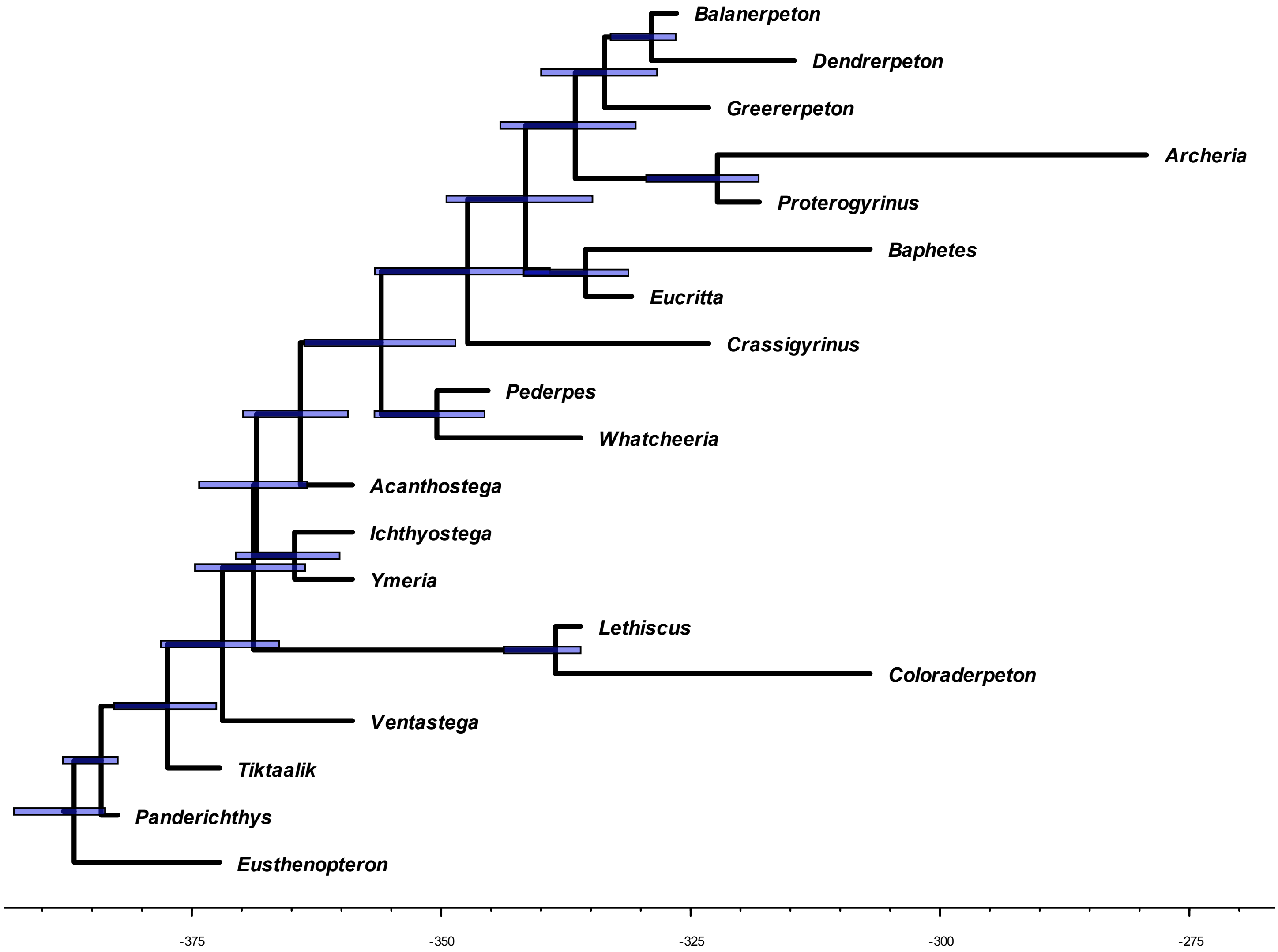


**Fig. 9.** The time-scaled phylogenetic tree inferred using Pardo et al.'s (2017) matrix. Node bars represent 95% highest posterior density (HPD) of node age estimates.

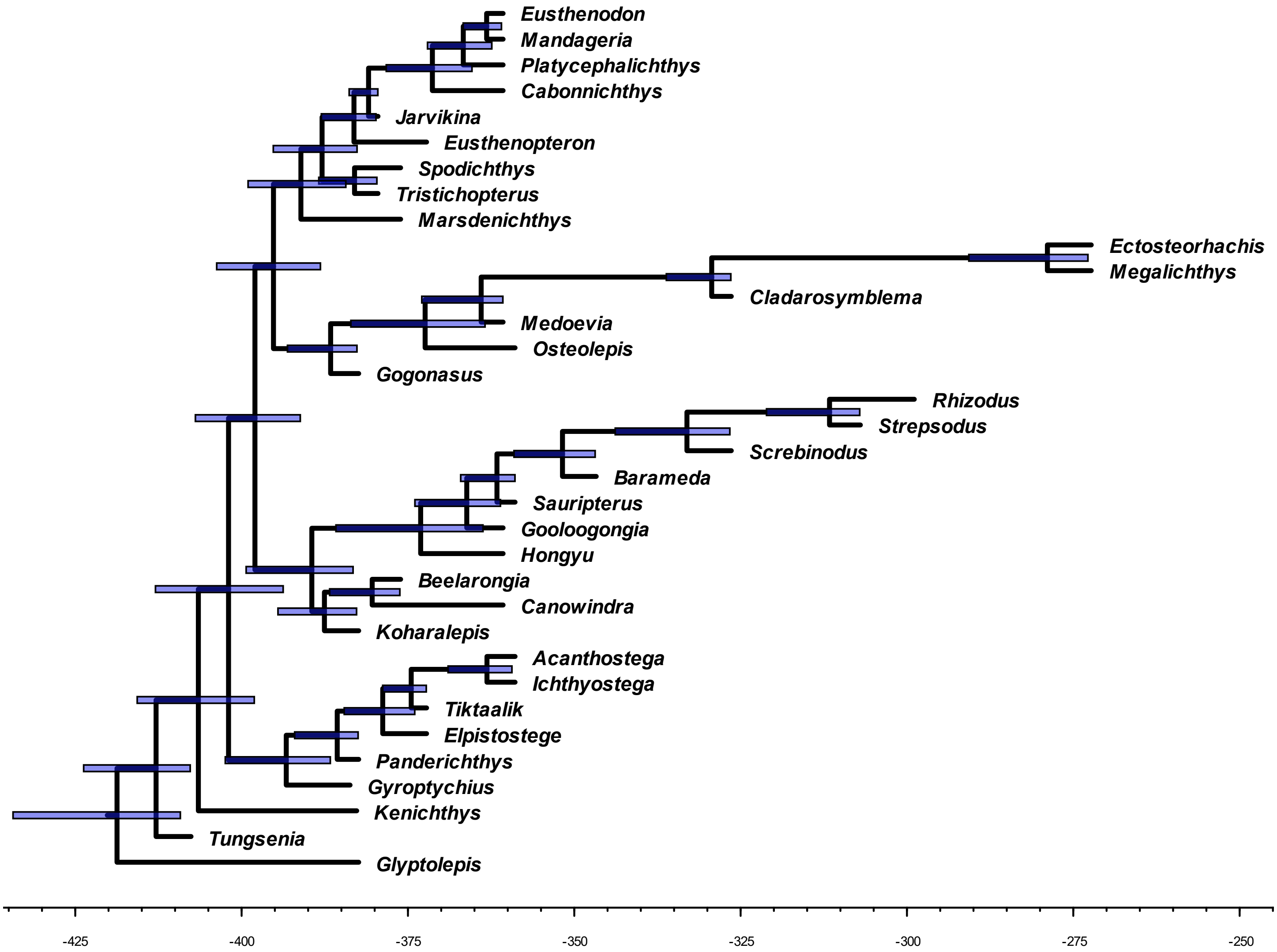


**Fig. 10.** The time-scaled phylogenetic tree inferred using Zhu et al.'s (2017) matrix. Node bars represent 95% highest posterior density (HPD) of node age estimates.

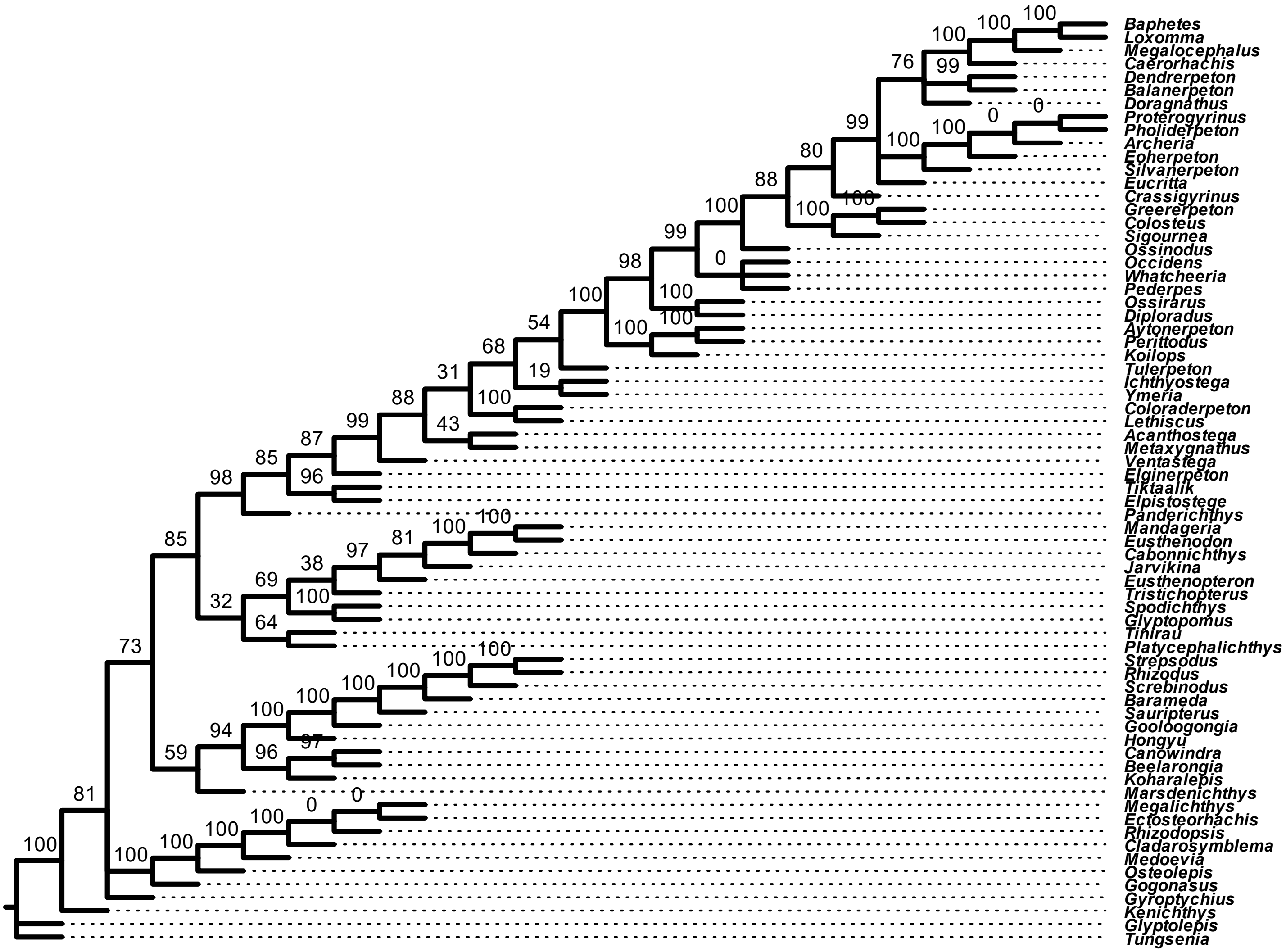


**Fig. 11.** Most of the supertree's confidence values at internal branches are above 50. However, there are some zeroes in regions that are otherwise well-supported in the source trees.

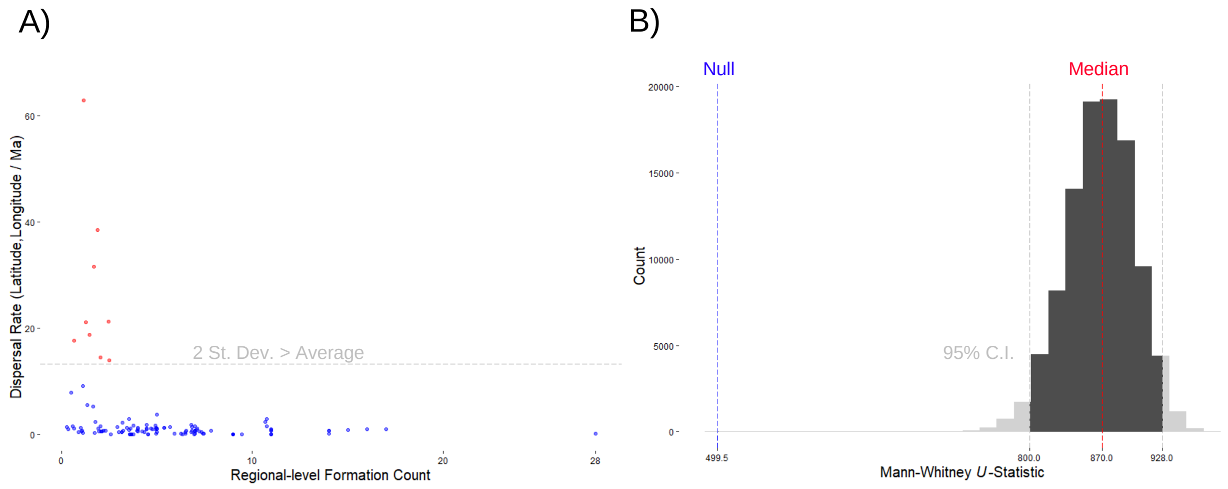


**Fig. 12. A)** Scatter-plot of the average dispersal rates over the regional-level formation counts for each branch of the phylogeny, using the Western Gondwana route scenario. Points colored by the dispersal rate being above or below two standard deviations greater than the average rate across the tree. **B)** Histogram of the bootstrapped *U*-statistics with values outside of the 95% confidence interval grayed out. The median and null expected *U*-statistics are indicated by the red and blue dotted lines, respectively. The null expected *U*-statistic is based on the null hypothesis that 50% of the regional-level formation counts with low dispersal rates will rank higher than formation counts with higher rates.

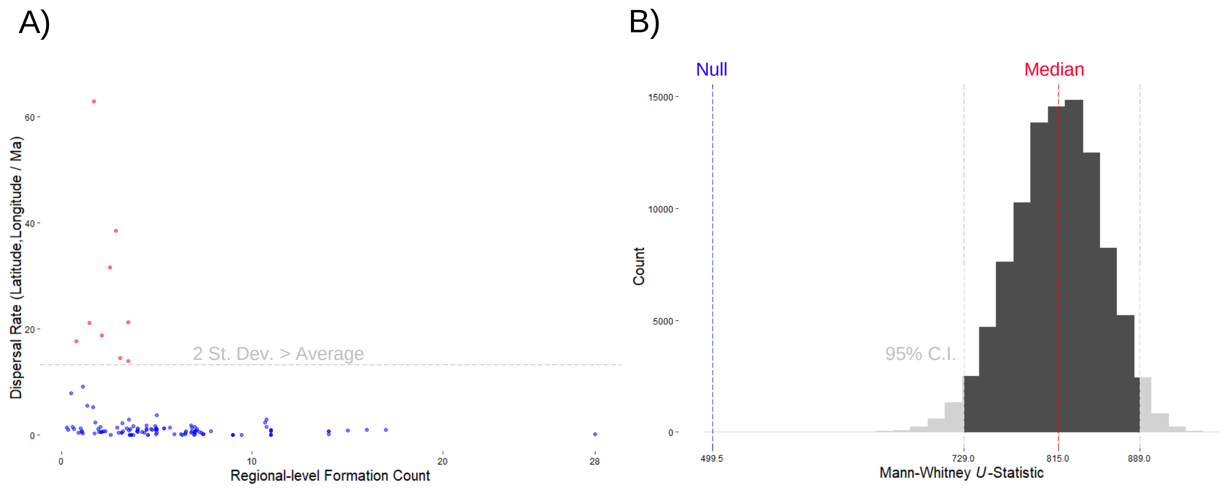


**Fig. 13. A)** Scatter-plot of the average dispersal rates over the regional-level formation counts for each branch of the phylogeny, using the direct route scenario. Points colored by the dispersal rate being above or below two standard deviations greater than the average rate across the tree. **B)** Histogram of the bootstrapped *U*-statistics with values outside of the 95% confidence interval grayed out. The median and null expected *U*-statistics are indicated by the red and blue dotted lines, respectively. The null expected *U*-statistic is based on the null hypothesis that 50% of the regional-level formation counts with low dispersal rates will rank higher than formation counts with higher rates.

**Supplementary files description**

*1. Supertree*

*1.1. Data*

- tree_2007_Friedman_etal.NWK
- tree_2012_Swartz.NWK
- tree_2017_Clack_etal.NWK
- tree_2017_Pardo_etal.NWK
- tree_2017_Zhu_etal.NWK

  We used these cladograms in Newick format when comparing the published source tree topologies with the supertree topology.

- tree_matrix_2007_Friedman_etal.NEX
- tree_matrix_2012_Swartz.NEX
- tree_matrix_2017_Clack_etal.NEX
- tree_matrix_2017_Pardo_etal.NEX
- tree_matrix_2017_Zhu_etal.NEX

  These NEXUS files contain data matrices and MrBayes blocks.

*1.2. Analysis*

- tree_mrbayes_output_2007_Friedman_etal
- tree_mrbayes_output_2012_Swartz
- tree_mrbayes_output_2017_Clack_etal
- tree_mrbayes_output_2017_Pardo_etal
- tree_mrbayes_output_2017_Zhu_etal

  These folders contain MrBayes outputs for each inference, including maximum clade credibility trees.

- tree_phydstar_supertree_rooted.trees

  These NEXUS files contain the final rooted supertree.

- tree_R_compare_trees.R

  We used the R script above to calculate normalized Robinson-Foulds distances between the supertree and the published source trees.

- tree_R_root_tree.R

  We used the R script above to root the supertree and save it in a NEXUS format.

- tree_sdm.txt

  We used the PHYLIP file above, which contains all five maximum clade credibility trees to build the distance supermatrix.

- tree_sdm_output_mat.txt

  This is the distance supermatrix.

*1.3. Publication*

- Gardner.etal.CRPalevol.Fig1.SupertreeFinal.v1.0.pptx
  We used Microsoft PowerPoint to prepare Figure 1.

- Gardner.etal.CRPalevol.Fig1.SupertreeRCode.v1.0.R
  We used the R script above to plot our supertree against the international geological time scale.

*2. Phylogeography*

*2.1. Data*

- Gardner.etal.FormationCounts.xlsx
  Regional- and stage-level formation counts from PBDB and Benton et al. (2013)

- Gardner.etal.GeoData.xlsx
  Paleocoordinate data entries from the PBDB.

*2.2. Analysis*

- Gardner.etal.VRCalc.xlsx
  Calculating dispersal rate for each branch of the phylogeny.

- Gardner.etal.VRRunComparisons.xlsx
  Comparing the ancestral states and rate scalars among the three independent variable rates runs.

- RegCount_WestGond_boot.R
  Bootstrapping Mann-Whitney U-test analyses using regional-level formation counts from Western Gondwana route scenario.

- RegCount_NorthEur_boot.R
  Bootstrapping Mann-Whitney U-test analyses using regional-level formation counts from Northern Euramerica route scenario.

- RegCount_Direct_boot.R
  Bootstrapping Mann-Whitney U-test analyses using regional-level formation counts from direct route scenario.
